# Disease-associated mutations in WDR34 lead to diverse impacts on the assembly and function of dynein-2

**DOI:** 10.1101/2022.03.31.486414

**Authors:** Caroline Shak, Laura Vuolo, Borhan Uddin, Yohei Katoh, Tom Brown, Aakash G. Mukhopadhyay, Kate Heesom, Anthony J. Roberts, Nicola Stevenson, Kazuhisa Nakayama, David J. Stephens

## Abstract

The primary cilium is a sensory organelle, receiving signals from the external environment and relaying them into the cell. Mutations in proteins required for transport in the primary cilium result in ciliopathies, a group of genetic disorders that commonly lead to the malformation of organs such as the kidney, liver and eyes and skeletal dysplasias. Motor proteins dynein-2 and kinesin-2 mediate retrograde and anterograde transport respectively in the cilium. WDR34, a dynein-2 intermediate chain, is required for the maintenance of cilia function. Here, we investigated *WDR34* mutations identified in Jeune syndrome, short-rib polydactyly syndrome or asphyxiating thoracic dysplasia patients. There is a poor correlation between genotype and phenotype in these cases making diagnosis and treatment highly complex. We set out to define the biological impacts on cilia formation and function of *WDR34* mutations by stably expressing the mutant proteins in WDR34 knockout cells. *WDR34* mutations led to different spectrums of phenotypes. Quantitative proteomics demonstrated changes in dynein-2 assembly, whereas initiation and extension of the axoneme, IFT-B protein localization, transition zone (TZ) integrity, and Hedgehog signalling were also affected.

**Summary statement:** Disease-associated mutations in WDR34 are found to have diverse impacts on ciliogenesis and cilia function following stable expression in a WDR34 knockout cell model.

## Introduction

The primary cilium is a microtubule-based structure that protrudes from the cell surface, its membrane continuous with the plasma membrane. The organelle acts as a cellular antenna, detecting stimuli from the surrounding environment and transmitting the information into the cell via receptors on its surface to mediate the appropriate response. Primary cilia have a major role in Hedgehog (Hh) signalling, directing proliferation, migration, and differentiation of cells with a particular importance during skeletal development (Goetz and Anderson, 2010). Defects in cilia assembly and function lead to a wide range of ciliopathies (Reiter and Leroux, 2017).

Assembly and ongoing function of the primary cilium is mediated by intraflagellar transport (IFT) of cargoes (Kozminski et al., 1993; Pigino, 2021; Webb et al., 2020). IFT is bidirectional and utilizes organelle specific motors kinesin-2 and cytoplasmic dynein-2 (hereafter referred to as dynein-2) to transit along microtubules doublets arranged along the central axis of the cilium (Pigino, 2021; Vuolo et al., 2020). Kinesin-2 works with IFT-B trains (Funabashi et al., 2018) to drive anterograde transport of cargoes along B-tubules of the microtubule doublets (Stepanek and Pigino, 2016) towards the tip of the cilium. Anterograde IFT trains also include IFT-A proteins and inactive dynein-2 (Jordan et al., 2018; Toropova et al., 2017; Toropova et al., 2019; Webb et al., 2020). At the tip, and likely at other sites along the axoneme {Nievergelt, 2022 #50}, reassembly of proteins and a change in conformation of dynein-2 leads to its activation, allowing retrograde trafficking of proteins by dynein-2 and IFT-A trains along the A-tubules of the microtubule doublets (Stepanek and Pigino, 2016). A ciliary necklace (Gilula and Satir, 1972) connects to the TZ, distal to the mother centriole, to form a diffusion barrier at the ciliary base from which proteins enter and exit the organelle. IFT is required both to maintain the TZ (Jensen et al., 2018; Vuolo et al., 2018) as well as enable crossing of this barrier (De-Castro et al., 2022; Park and Leroux, 2022).

Metazoan dynein-2 is formed of multiple subunits (Asante et al., 2014; Vuolo et al., 2020) built around two copies of the dynein heavy chains ((encoded by *DYNC2H1*) (Pazour et al., 1999; Porter et al., 1999; Signor et al., 1999)). The motor complex is asymmetric, containing two different dynein-2 intermediate chains (Asante et al., 2014; Toropova et al., 2019): DYNC2I1 (originally identified as FAP163 in *Chlamydomonas* (Patel-King et al., 2013) and commonly called WDR60 in mammals) and DYNC2I2 (originally identified as FAP133 in *Chlamydomonas* (Rompolas et al., 2007) and commonly called WDR34 in mammals). They each contain N-terminal regions that bind dynein light chains (Pazour et al., 1998) and a C-terminal WD-propellor structure, each of which binds one dynein-2 heavy chain (Toropova et al., 2019). A dimeric dynein-2-specific light intermediate chain, commonly called LIC3 (encoded by *DYNC2LI1* (Grissom et al., 2002; Hou et al., 2004)), in turn binds to and stabilizes the individual heavy chains. Light chains DYNLL1 and DYNLL2 (LC8 family), DYNLRB1 and DYNLRB2 (roadblock family), DYNLT1 and DYNLT2 (Tctex family) are shared with dynein-1; *DYNLT2B* encoding TCTEX1D2 is unique to dynein-2 (Asante et al., 2014). We use the more common names throughout for ease of reading.

Mutations in each of the genes encoding dynein-2 subunits lead to ciliopathies, specifically, those associated with skeletal abnormalities (Huber and Cormier-Daire, 2012). In particular, multiple studies have identified *WDR34* mutations in individuals with Jeune syndrome or Short-rib polydactyly syndrome (SRPS) (Huber et al., 2013; Schmidts et al., 2013; You et al., 2017) with more continuing to be identified (Yin et al., 2020). Clinical phenotypes differ between individuals (Schmidts et al., 2013) but all have skeletal defects in common. Supplementary Table 1 shows the diversity of clinical outcomes for individuals with mutations in *WDR34* emphasising the deficit in correlation between genotype and phenotype (Huber et al., 2013; Schmidts et al., 2013; You et al., 2017). Supplementary Table 2 shows predictions of the functional impact of mutations in WDR34 again highlighting the lack of clear correlation between genotype and phenotype.

Although these mutations are often homozygous, they are not always found in isolation (Schmidts et al., 2013). This is also true of other ciliopathies for example where mutations in *TTC21B* (which encodes IFT139) can cause both isolated nephronophthisis and syndromic Jeune asphyxiating thoracic dystrophy as well as showing extensive genetic interactions with other ciliopathy loci (Davis et al., 2011). It has been suggested that mutations could act in trans and digenic inheritance has been proposed in some cases, complicating the explanation of clinical presentations. For example, patients with p.A22V mutations in WDR34 were also heterozygous for other mutations in *WDR19* and *IFT140* that are predicted to be deleterious to function (Schmidts et al., 2013) and Supplementary Table 2.

Knockdown and knockout (KO) models of WDR34 have been studied in cells and model organisms (Huber et al., 2013; Schmidts et al., 2013; Wu et al., 2017). A WDR34 KO cell culture model containing only the first 24 or 44 residues of WDR34 resulted in longer cilia, mis-localization of the IFT-B protein IFT88 and impaired Hh signalling (Tsurumi et al., 2019). Similar but more pronounced phenotypes were observed in WDR60 KO cells (Hamada et al., 2018; Tsurumi et al., 2019, Vuolo et al., 2018). Different WDR34 KO cell lines (Tsurumi et al., 2019; Vuolo et al., 2018) and a KO mouse model (Wu et al., 2017) have been reported to have very short primary cilia; notably these retain expression of a longer N-terminal WDR34 region of 114-146 amino acids. These KO models, show changes in IFT88 and IFT140 distribution (Vuolo et al., 2018; Wu et al., 2017), impaired Hh signalling (Wu et al., 2017), and reduced assembly of the dynein-2 holocomplex (Vuolo et al., 2018).

In this study we set out to determine the impact solely of the mutations in WDR34 to gain a better understanding of the disease phenotypes as well as the role of this key dynein-2 subunit. Here, we have examined the ability of clinically identified point mutations in WDR34 to restore normal ciliary structure and function to *WDR34* KO cells. In order to achieve this, we have reconciled previous work using different *WDR34* KO models and then expressed *WDR34* mutants in a null background. Using proteomics, biochemistry, and cellular imaging, our data show that WDR34 mutations have diverse impacts on dynein-2 integrity, ciliogenesis, cilia length control, and signalling. Notably, many mutations can restore significant aspects of cilia function and selective defects are seen with each mutation.

## Results

Several different WDR34 KO models have been generated with some key differences in the observed phenotypes. Experiments in mouse embryonic fibroblasts and retinal pigment epithelial (RPE-1) cells introduced frameshift mutations into the *WDR34* gene (Figure 1A) (Vuolo et al., 2018; Wu et al., 2017) that caused a severe reduction in both the frequency of ciliated cells and cilia length. These mutants retain expression of an N-terminal polypeptide of WDR34. In contrast, other WDR34 KO cells that remove expression completely have longer cilia than the WT RPE1 line (Tsurumi et al., 2019). To reconcile these findings, Tsurumi et al. (2019) provided evidence that the residual N-terminal WDR34 fragment may exert a dominant negative effect on cilia formation by binding to and sequestering the dynein light chains DYNLL1 and DYNLL2 (depicted in Figure 1A). WDR34 KO cell lines generated with p.Val119Alafs*4 or p.Thr151Thrfs*41 mutations and encoding similar N-terminal regions also assemble short cilia (Tsurumi et al., 2019). We considered that the same could also be true for WDR34 for the disease-associated mutation p.Q158*.

**Figure 1.**
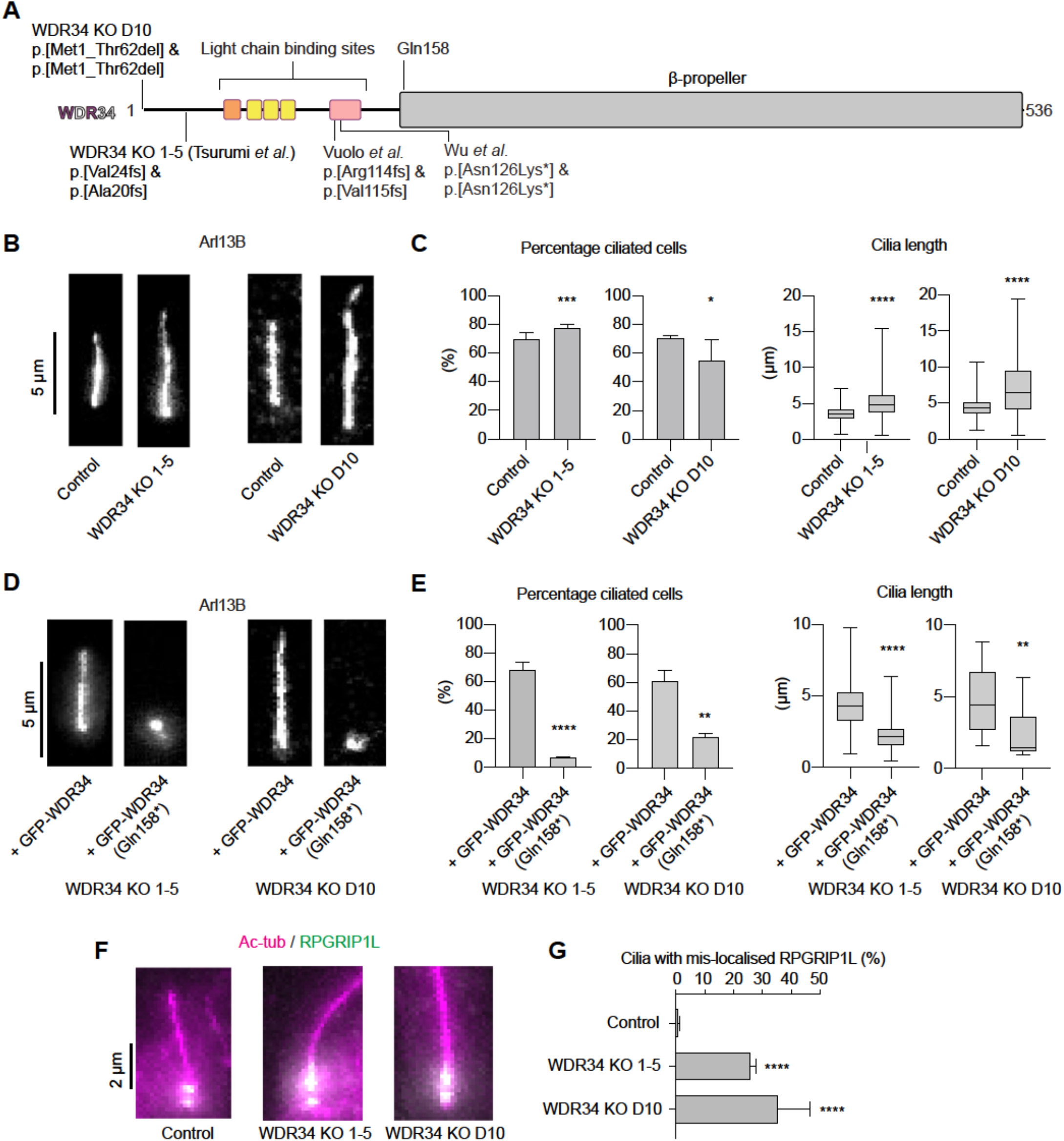
WDR34 KOs in RPE-1 cells build a cilium albeit with TZ defects. (A) Domain architecture of human WDR34 highlighting regions targeted for genome editing in indicated studies. Location of patient mutation WDR34 Gln158* is also denoted. (B) Representative images of WDR34 KO cilia with respective matched control in RPE-1 cells. Cells were fixed and stained with Arl13B to identify cilia. (C) Percentage of ciliated cells and cilia length of WDR34 KO cells with respective matched control in RPE-1 cells. For percent ciliated cells, 1275 control, 1327 WDR34 KO 1-5 and 446 control, 680 WDR34 KO D10 cells were quantified. Values are mean and SEM. For cilia length, 890 control, 1027 WDR34 KO 1-5; and 341 control, 429 WDR34 D10 cilia were measured. Cilia length of WDR34 KOs were increased in comparison to matched control, * p < 0.05, p < 0.001, **** p < 0.0001 (Mann-Whitney test). Whiskers represent minimum and maximum values. Three independent experiments were performed. (D) Representative images of WDR34 KO 1-5 and D10 cilia with expression of either GFP-WDR34 or GFP-WDR34 Gln158*. Cells were fixed and stained with Arl13B to identify cilia. (E) Percentage of ciliated cells and cilia length of WDR34 KO 1-5 and D10 cell lines with expression of either GFP-WDR34 (WDR34) or GFP-WDR34 p.Q158* (p.Q158*). GFP constructs were stably expressed in WDR34 KO 1-5 and in WDR34 KO D10 cell line. Expression of p.Q158* in WDR34 KOs led to severe reduction in frequency and cilium length. Asterisks indicate p-values for Mann-Whitney test. **p < 0.01, ****p < 0.0001. For percent ciliated cells, 1070 (WDR34 KO 1-5 + WDR34), 1091(WDR34 KO 1-5 + p.Q158*) were quantified. For WDR34 KO D10 +GFP-WDR34 and WDR34 KO D10 +GFP-WDR34 p.Q158*, 57 and 56 GFP+ cells were quantified for percent ciliated cells. Values are mean and SEM. For cilia length, 739 (WDR34 KO 1-5 + WDR34), 81 (WDR34 KO 1-5 + p.Q158*), 24 (WDR34 KO D10 + WDR34), 8 (WDR34 KO D10 + p.Q158*) cilia were measured. Whiskers represent minimum and maximum values. Three independent experiments were performed. (F) Representative images of control, WDR34 KO 1-5 and D10 TZ in RPE-1 cells. Cells were fixed and stained with acetylated tubulin (magenta) to identify cilia and RPGRIP1L (green) as a TZ marker. (G) Percentage cilia with RPGRIP1L extending into the cilium. 230 WT, 276 WDR34 KO D10, 245 WDR34 KO 1-5 cilia were analysed. Significant increase in percentage cilia with RPGRIP1L mis-localised in WDR34 KOs vs control (p < 0.0001, One-way ANOVA followed by Kruskal-Wallis test). Values are mean and SEM. Three independent experiments were performed.

### Variation of cilia phenotypes in different WDR34 KO cell lines

To validate these findings, we generated a new RPE-1 WDR34 KO cell line in which the start codon of *WDR34* was deleted (termed here WDR34 KO D10) (Supplementary Figure S1) and compared this to the RPE-1 WDR34 KO 1-5 cell line generated by Tsurumi et al. (2019), WDR34 KO 1-5. Cells were serum starved to allow analysis of cilia. In both cases, the WDR34 KO cells were able to form cilia with a frequency comparable to control cells, with a minor increase in cilia length (Figure 1B, C).

### WDR34 p.Q158* inhibits ciliogenesis

These data support the notion that N-terminal WDR34 fragments can cause a more severe ciliary phenotype than if the entire WDR34 protein is absent (Tsurumi et al., 2019). To test if the p.Q158* mutant could exert an inhibitory effect on cilia formation, we expressed either GFP-tagged WDR34 or GFP-tagged WDR34-p.Q158* in WDR34 1-5 KO RPE-1 cells. Whereas cilia formation was not affected by expression of full-length WDR34, expression of WDR34 p.Q158* in WDR34 KO cells resulted in a severe decrease in both the fraction of ciliated cells and cilia length (Figure 1D, E and Supplementary Figure S1C). Notably, this was not the case when p.Q158* was expressed in a WT background (Supplementary Figure S1C). Proteomics experiments demonstrated that WDR34 p.Q158* retained the ability to bind to dynein light chain DYNLL1 and other dynein-2 subunits, consistent with a sequestering effect (Supplementary Table S3).

### TZ impairments in WDR34 KO lines D10 and 1-5

At the base of the cilium, the TZ acts as a gatekeeper between the ciliary compartment and the cytoplasm, helping to determine the protein composition of the cilium (Shi et al., 2017). Previous studies demonstrated that loss of WDR60 is associated with mis-localisation of the TZ marker RPGRIP1L (Vuolo et al., 2018). We therefore tested the localisation of RPGRIP1L in RPE-1 WDR34 KO cells. Whereas RPGRIP1L was restricted to a region near the ciliary base in control cells, in both WDR34 KO D10 and WDR34 KO 1-5 cell lines, RPGRIP1L was mis-localised and extended into the cilium in a significant fraction of cells (Figure 1F, G). We conclude that while WDR34 is not critical for cilium formation in RPE-1 cells, consistent with the findings of Tsurumi et al., (2019), but is required for proper organisation of the TZ as is also true for WDR60 (Vuolo et al., 2018).

### Analysis of disease-associated mutations in *WDR34*

Twelve point mutations and one truncation mutation in *WDR34* were investigated (Figure 2A). Eleven of the residues are well conserved amongst species (summarized in Supplementary Figure S3), suggesting a significant role in protein function. Figure 2B shows a schematic of the location of individual subunits including WDR34 within the dynein-2 complex. The location of each mutant relative to its position within the dynein-2 complex is shown in Supplementary Figure S4. We include predictions of the potential functional impact of these mutations from PolyPhen2 (Adzhubei et al., 2010), SIFT (Ng and Henikoff, 2003), and PROVEAN (Choi et al., 2012) in Supplementary Table 2. To study these WDR34 mutations, we used the WDR34 KO cell line 1-5 (Tsurumi et al., 2019) cells and transduced them with lentiviruses to generate polyclonal cell lines stably expressing wild-type (WT) GFP-WDR34, specific WDR34 mutants, or GFP only. All constructs were expressed as intact proteins at the expected molecular weight, as shown by immunoblotting of cell lysates, but at different levels (Figure 2C and Supplementary Figure S2A). Notably, GFP-WDR34-p.Q158* was expressed at a low level. Varying phenotypes in cells expressing WDR34 mutations may result from changes in protein function but may also be due to the different protein levels. Therefore we also analysed the relative stability of each mutant using cycloheximide (to inhibit further protein synthesis) and MG132 (to inhibit proteasomal degradation). Figure 2C shows that p.A22V, pQ158*, p.R206C, and p.T354M have similar stability to WT WDR34. In contrast, p.C148F, p.R182W, p.P390L, p.G393S, p.S410I, p.R447Q, and p.R447W all appear less stable (enhanced degradation with cycloheximide). WDR34 p.A341V is expressed at low levels making interpretation difficult. Notably, p.K436R shows enhanced stability compared to both WT and other WDR34 mutants. p.R447W appears more susceptible to degradation than p.R447Q, even in the presence of MG132.Full gels are shown in Supplementary Figure 2B along with tubulin as a control.

**Figure 2.**
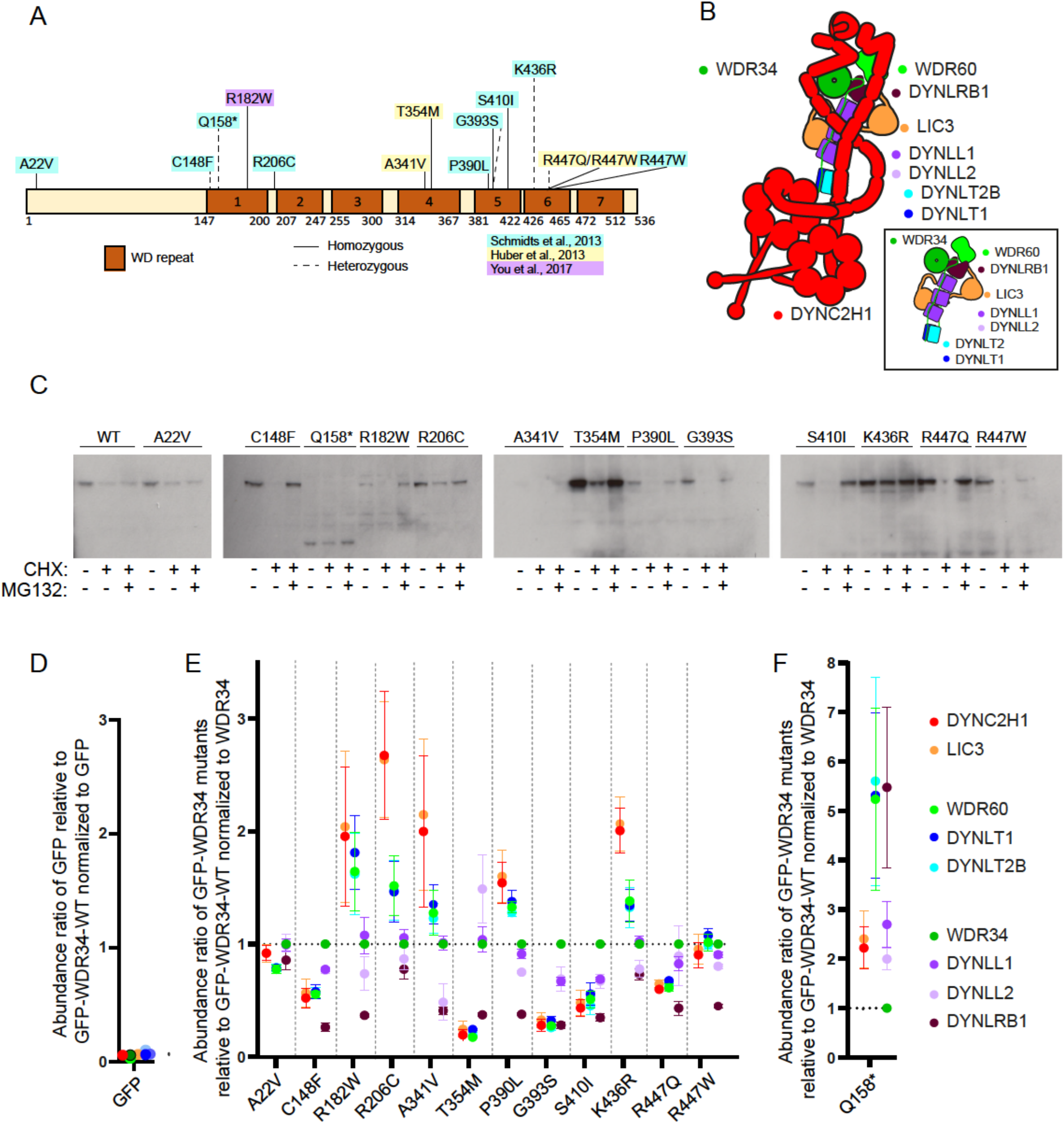
WDR34 mutations in patients. (A) Whole exome screening identified 15 point mutations located throughout the length of the protein. The 12 missense mutations and one resulting in a pre-mature stop codon are shown. (B) Schematic of dynein-2 (adapted from (Vuolo et al., 2020)). Colouring is the same as in subsequent plots. Note there remains some uncertainty on the light chain subunit composition regarding DYNLL1 and DYNLL2 as well as DYNLT1 and DYNLT2B. (C) Mutants were analysed by immunoblotting to determine relative expression level (see also Supplementary Figure 2A) and relative stability by treating cells with cycloheximide or cycloheximide plus MG132 for 16 h prior to lysis and immunoblotting. (D) Plot showing relative abundance of dynein-2 subunits interacting with GFP only. These data compare cells expressing GFP with those expressing GFP-WDR34-WT normalised to GFP abundance. (E) WDR34 mutants interact differently with DYNC2H1-LIC3 and WDR60-DYNLT2B-DYNLT1 subcomplexes, and DYNLRB1. RPE1 WDR34 KO cell lines expressing GFP proteins were serum-starved for 24 h to induce ciliogenesis and harvested to purify GFP proteins by GFP-trap. Proteomics of GFP-WDR34 proteins by TMT-labeling was used to relatively quantify interaction with other dynein-2 proteins. Protein abundances were normalized to WDR34 and compared to the GFP-WDR34-WT sample to calculate protein abundance ratios. (F) Separate plot for GFP-WDR34-p.Q158* plotted along with GFP-WDR34 WT (as in E) owing to different scaling of the y-axis.

### Impact of WDR34 mutations on dynein-2 complex assembly

Figure 2B shows a schematic of the location of dynein-2 subunits colour-coded as in the subsequent plots. We utilized 16-plex multiplex tandem-mass-tag (TMT)-labelling to relatively quantify the interaction of WDR34 point mutant proteins with other dynein-2 components. To prepare cell samples for proteomics, clarified protein lysates were incubated with GFP-Trap beads to identify proteins interacting with GFP-WDR34. Purified proteins were labelled with tandem labelled tags prior to MS analysis to allow the relative quantification of proteins in each sample. Figure 2D shows that we do not efficiently detect any proteins in GFP-nanotrap pull downs from cells expressing GFP only relative to those expressing GFP-WDR34-WT. These data are normalized to the abundance of GFP. Figure 2E, 2F show data from cells expressing GFP-WDR34 mutants relative to GFP-WDR34-WT; these data are normalized to the abundance of WDR34 to counter differential expression levels and stability of each mutant. Dynein-2 complex components DYNC2H1, WDR60, LIC3, DYNLL1, DYNLL2, DYNLRB1, DYNLT1 and DYNLT2B were identified as interacting partners across all samples in all three experimental repeats (Figure 2E, F). DYNLRB2 and DYNLT3 were not identified in these experiments. These data are summarized graphically in Supplementary Figure S5 where the central circle represents the abundance relative to GFP-WDR34-WT. Those subunits shown within the circle are identified more readily suggesting a tighter interactions, those outside the circle indicate less abundance and likely weaker interactions.

Dynein-2 can be split into three subcomplexes (Hamada et al., 2018), DYNC2H1-LIC3, WDR60-TCTEX1D2-DYNLT1, and WDR34-DYNLL1/DYNLL2-DYNLRB1. Association of WDR34 A22V, R447Q and R447W with DYNC2H1-LIC3 and WDR60-TCTEX1D2-DYNLT1 were similar to WDR34 WT. Mutations that resulted in a >50% reduction in interaction were C148F, T354M, G393S and S410I, whereas mutations that resulted in a 2-fold increased interaction with DYNC2H1-LIC3 were p.Q158*, p.R182W, p.R206C, p.A341V and p.K436R. These WDR34 mutants also resulted in a modestly increased interaction with WDR60-DYNLT2B-DYNLT1 (Figure 2E,F).

Formation of the third dynein-2 subcomplex, WDR34-DYNLL1/DYNLL2-DYNLRB1 involves binding of the light chains to the N-terminal region of WDR34 (Toropova et al., 2019). Interestingly, apart from WDR34-K436R, interaction with DYNLRB1 was reduced by more than half in all WDR34 point mutants located in a WD domain, suggesting defective assembly of DYNLRB1 in dynein-2 may be a common phenotype in WDR34-related disorders. The association of DYNLL2 with WDR34 was only significantly reduced in cells expressing WDR34 p.A341V, whereas DYNLL1 was not markedly affected. The differences in light chain binding to WDR34 mutants suggest they bind independent of one another. These results show that WDR34 mutations impact dynein-2 assembly by either enhancing or decreasing association depending on the mutation.

### Impact on ciliogenesis and cilia length

All GFP-WDR34 proteins showed a significant cytosolic pool (Figure 3A) with some minor localization to centrioles. Consistent with immunoblotting, GFP-WDR34-p.Q158* was expressed at a lower level and was more prominently observed at the ciliary base. To examine the effect of WDR34 mutations on the regulation of ciliogenesis, cells were immunolabelled to detect the cilia membrane marker ARL13B (Figure 3A). We scored the percentage of ciliated cells and cilia length, testing all cell lines against the parental WDR34 KO expressing GFP only, experiments included WT RPE1 cells for comparison.

**Figure 3.**
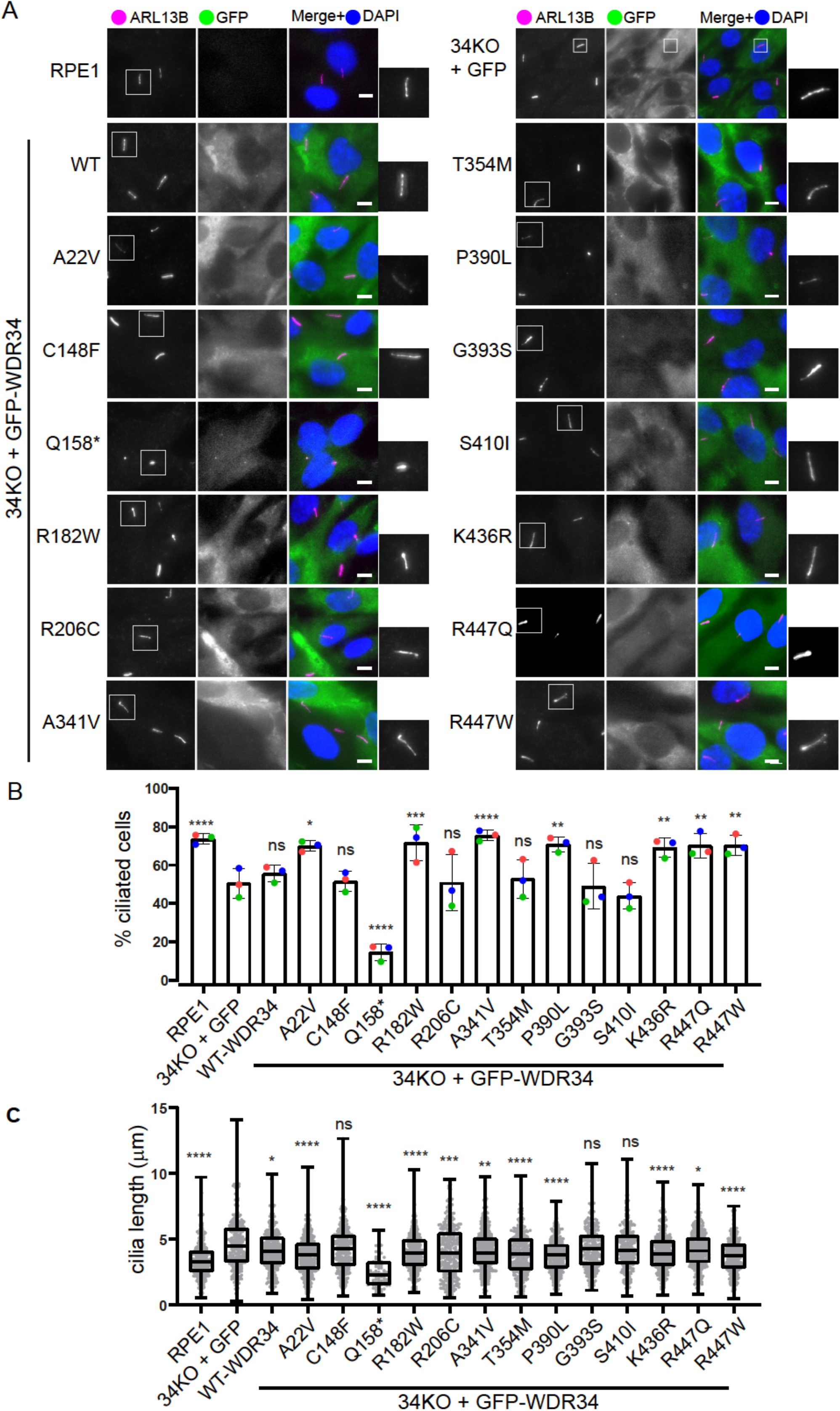
Formation and length of cilia is affected by WDR34 mutations. (A) WT RPE1 and WDR34 KO cell lines expressing GFP proteins (green) were serum-starved for 24 h to induce ciliogenesis, fixed with −20°C methanol and labelled with an antibody to detect ARL13B (magenta). Scale bar = 5 μm. Two-fold enlargements of cilia staining (ARL13B) are shown. (B) Percentage of ciliated cells in each cell line. Data pooled from three independent experiments with between 455 and 677 cells quantified. Data show means and standard deviation, n=3. *P=<0.05 using Tukey’s multiple comparisons test. (C) Cilium length in each cell line. Data pooled from three independent experiments with between 78 and 406 cilia quantified. Box and whisker plot shows the median, interquartile ranges, minimum and maximum values, n=3. *P=<0.05 using Tukey’s multiple comparisons test.

Expression of some mutants in the WDR34 KO line (p.A22V, p.R182W, p.A341V, p.P390L, p.K436R, p.R447Q, and p.R447W) led to an increase in the percentage of ciliated cells. The truncated version, p.Q158*, however, led to a significant reduction in the ability of cells to form cilia, consistent with a dominant negative impact on ciliogenesis (see Figure 1 and (Tsurumi et al., 2019)). The increase in cilia length in WDR34 KO cells is reversed in cells expressing WT WDR34 and almost all mutant proteins. The p.C148F, p.G393S and p.S410I mutations were not able to rescue cilia length. The p.Q158* mutation led to further reduction in cilia length, again suggesting the fragment acquired functions that are restrictive to cilia assembly (Tsurumi et al., 2019).

To summarize the proteomics and ARL13B data, WDR34 mutants that significantly increase the proportion of ciliated cells either showed no change or enhanced dynein-2 assembly. Conversely, mutations unable to rescue cilia length also showed reduced dynein-2 assembly.

### Impact on the localization of RPGRIP1L

The TZ at the ciliary base mediates the entry and exit of proteins from the cilium. This diffusion barrier has been found to be disrupted in WDR60 KO cells (Vuolo et al., 2018). We immunolabelled ciliated cells with an antibody against RPGRIP1L, a TZ protein, to investigate whether this defect also arises in WDR34 KO cells and if so whether it could be “rescued” by any of the disease-associated mutations in WDR34 (Figure 4A). As expected in RPE1 WT cells, RPGRIP1L was localized at the ciliary base as compact circular dots, adjacent to the basal body. In the WDR34KO-GFP cells, a proportion of the cells had RPGRIP1L staining which stretched and extended towards the cilium, similar to WDR60 KO cells (Vuolo et al., 2018). This was significantly reduced upon expression of WT WDR34. RPGRIP1L localization was similarly rescued by nearly all mutations, including the p.Q158* truncation (Figure 4B), the only exception being the p.R447Q mutant. This supports a role for WDR34 in maintaining TZ integrity. Notably, of p.R447Q and p.R447W, only p.R447Q fails to restore the localization of RPGRIP1L. Our proteomics study showed that the interaction of WDR34-p.R447Q, but not p.R447W, with other dynein-2 proteins is slightly decreased.

**Figure 4.**
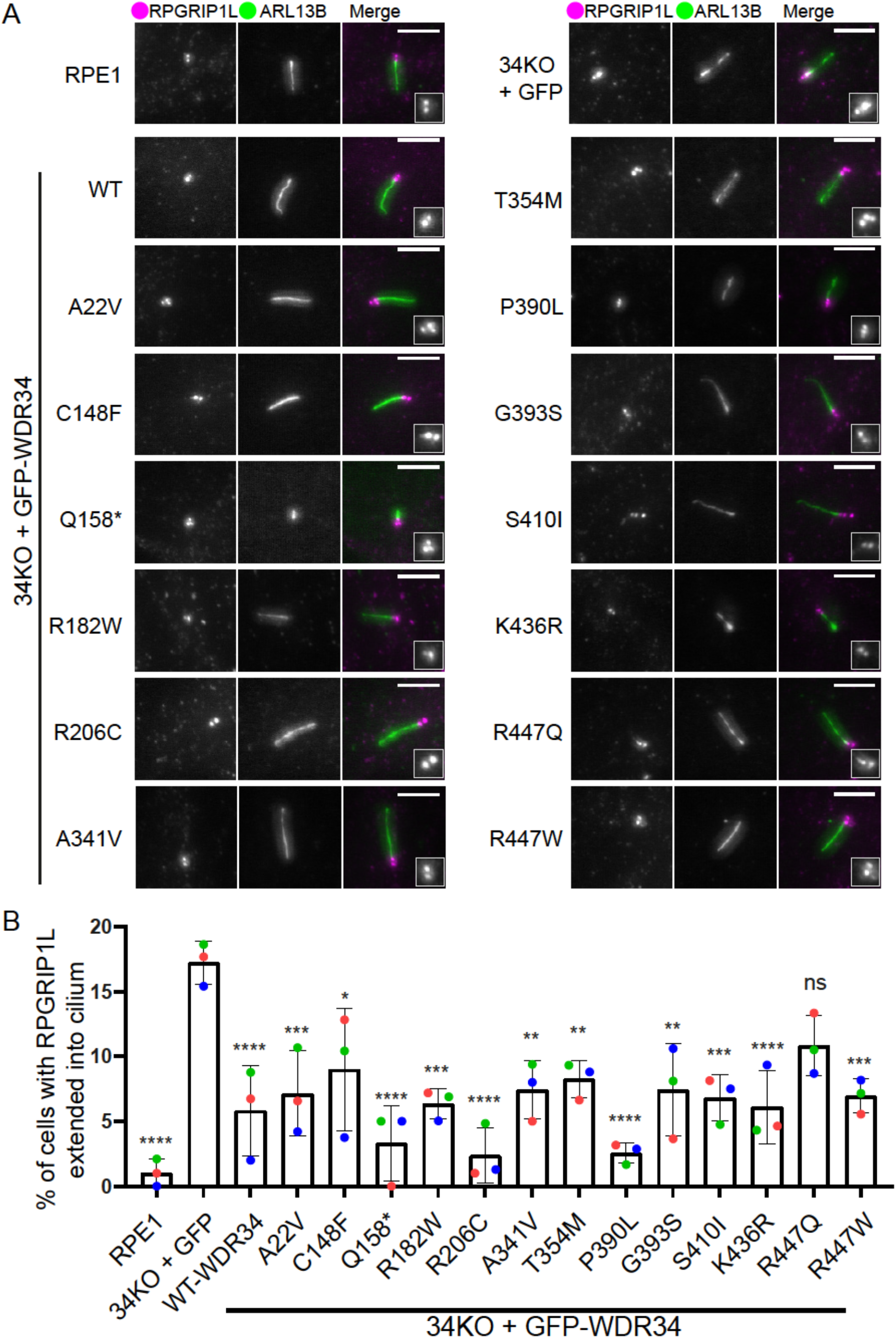
RPGRIP1L localization is only affected by WDR34-p.R447Q. (A) WT RPE1 and WDR34 KO cell lines expressing GFP proteins were serum-starved for 24 h to induce ciliogenesis, fixed with −20°C methanol and labelled with antibodies to detect ARL13B (green) and RPGRIP1L (magenta). Scale bar = 5 μm. 1.5-fold enlargements of RPGRIP1L staining are shown. (B) Percentage of cells with RPGRIP1L extension into the cilia. Between 95 and 455 cilia quantified in each case. Data show means and standard deviation, n=3 independent experiments. *P=<0.05 using Tukey’s multiple comparisons test.

### IFT140 localization

To analyse the effects of the WDR34 mutations on IFT-A proteins, IFT140 was immunolabelled alongside ARL13B as a ciliary marker (Figure 5). In WDR34 KO cells, IFT140 localizes to the ciliary base, as it does in WT RPE-1 cells. The specific lot of IFT140 antibody used also stains non-specific nuclear dots, as observed by others (Hamada et al., 2018; Hirano et al., 2017; Takahara et al., 2019), and described by the manufacturer. This complicates analysis and so we have not applied automated quantification to these data but instead scored for the robust presence of IFT140 by eye. Specific IFT140 staining was observed at the ciliary base in WDR34 KO cells expressing GFP only and nearly all WDR34 point mutations. This shows that WDR34 KO cells retain sufficient retrograde IFT function to maintain IFT140, and by inference the IFT-A complex, at the base. The only exception to this is seen with p.Q158* cells where IFT140 was seen throughout the length of the cilium, again suggesting that this truncated WDR34 protein results in dominant negative functions within the cilium, here in the assembly or localization of IFT-A.

**Figure 5.**
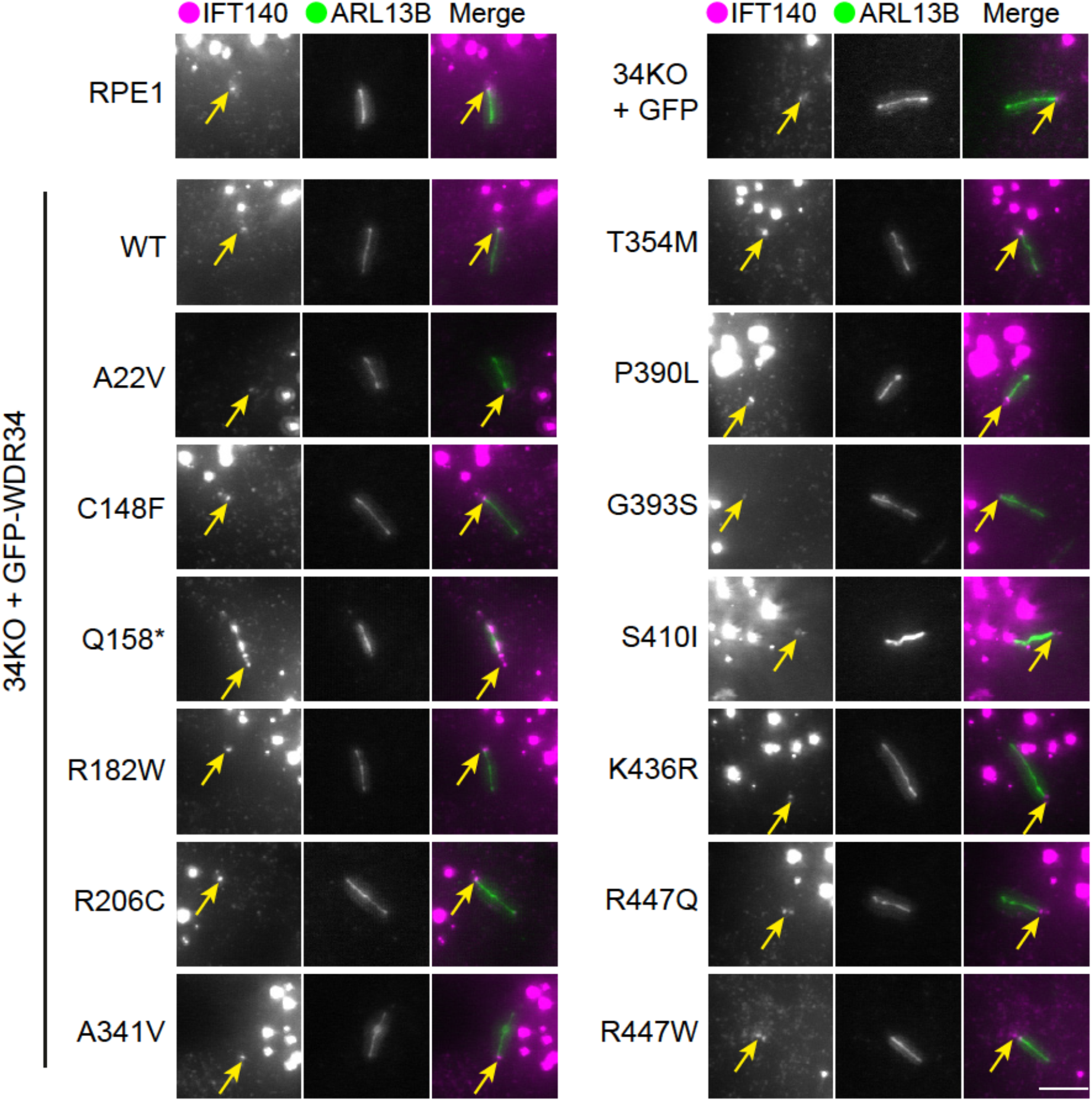
IFT140 localization is not affected by WDR34 missense mutations. WT RPE1 and WDR34 KO cell lines expressing GFP proteins were serum-starved for 24 h to induce ciliogenesis, fixed with −20°C methanol and labelled with antibodies to detect ARL13B (green) and IFT140 (magenta) antibodies. Scale bar = 5 μm.

### Accumulation of IFT88 within the cilium

We next investigated the localization of the IFT-B protein, IFT88, as a well-characterized reporter of IFT function as it accumulates at ciliary tips when retrograde defects occur. IFT88 is localized at the ciliary base in wildtype cells, but at the tips in WDR60 and WDR34 KO cells (Vuolo et al., 2018). We classified the distribution of IFT88 (Figure 6A) into groups: base only, base and tip, throughout, and uniformly throughout (Figure 6B). In RPE1 cells, IFT88 was mostly observed at the ciliary base and tip, or at the base only. In WDR34KO-GFP cells, the “base and throughout” localization became the most dominant distribution, whereas WDR34 WT rescued IFT88 distribution. The p.Q158* truncation results in a uniform distribution of IFT88 throughout the cilia. WDR34-p.A341V and p.T354M expressing cells also had a small increased proportion of cilia with IFT88 along the cilium. This change is concurrent with both a decrease in proportion of cells with IFT88 at the base only or base and tip, suggesting both localizations reflect normal IFT-B function.

**Figure 6.**
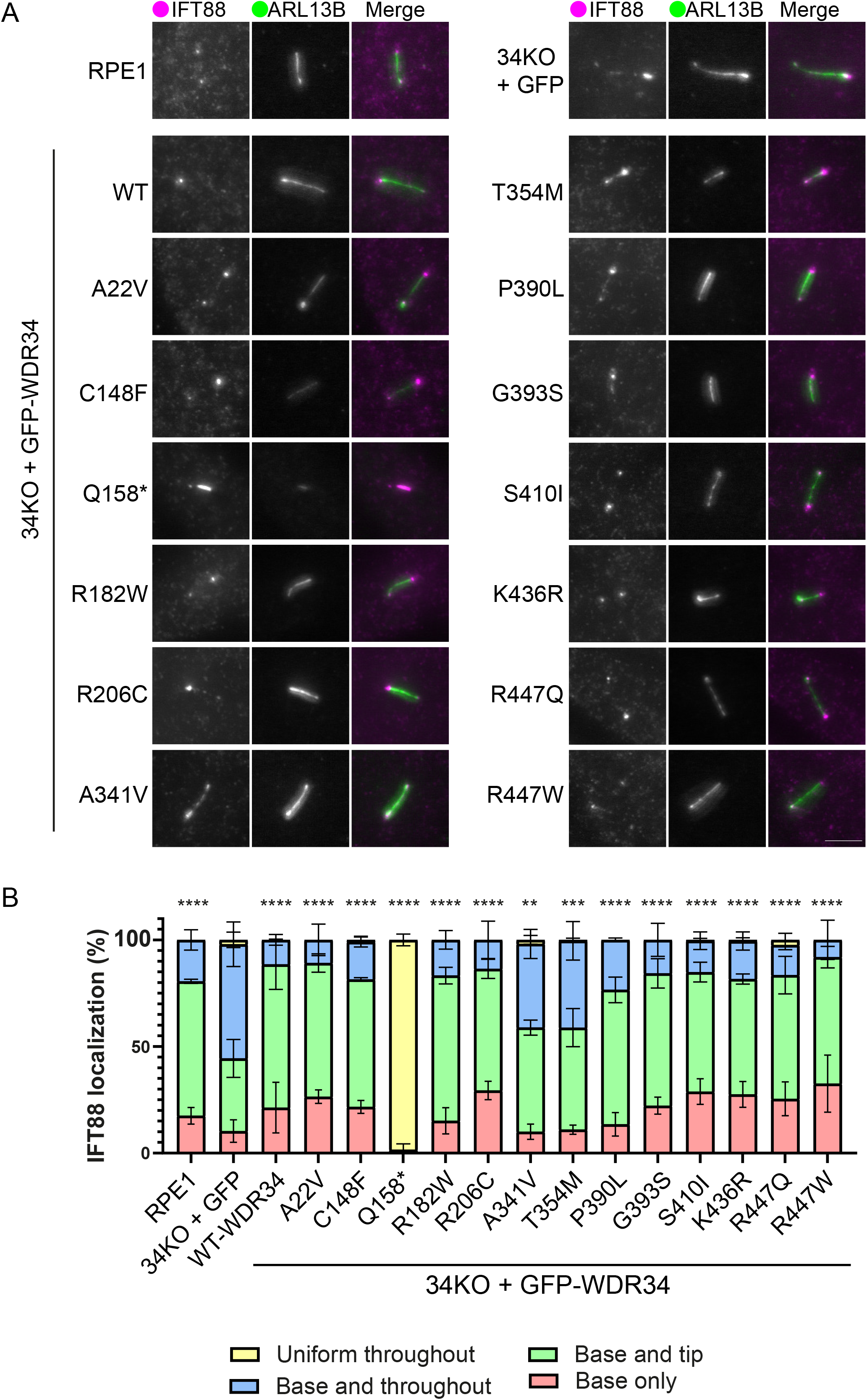
IFT188 localization is only affected by WDR34-p.A341V and WDR34-p.T354M. (A) WT RPE1 and WDR34 KO cell lines expressing GFP proteins were serum-starved for 24 h to induce ciliogenesis, fixed with −20°C methanol and labelled with antibodies to detect ARL13B (green) and IFT88 (magenta) antibodies. Scale bar = 5 μm. (B) Percentage of cells with various IFT88 distribution in each cell line. Cells were scored for IFT88 localization at the base only, throughout the cilium only, at the base and tip but not throughout the cilium, or at both base and cilia. Between 102 and 433 cilia quantified. Data shows mean and standard deviation, n=3 independent experiments. For statistical analysis, “uniform throughout” with “base and throughout”, and “base and tip” with “base only” were grouped together to compare for improper IFT88 localization. *p=<0.05 using Tukey’s multiple comparisons test.

### Basal levels and SAG-stimulated localization of Smoothened in the cilium

The studies shown above provide insight to the assembly of the dynein-2 and IFT complexes. It is, however, the functional changes resulting from defects in IFT, such as trafficking of receptors and signalling proteins, that translate these mutations into disease phenotypes. Since WDR34 mutations lead to skeletal ciliopathies, Hh signalling is of key interest owing to its well-established roles in skeletal development. In cell culture, Hh signalling can be artificially stimulated with smoothened agonist (SAG). Under basal conditions, the GPCR family member smoothened (SMO) is typically excluded from cilia at the basal state and SAG stimulation promotes its entry into and accumulation within cilia.

As expected in basal conditions, SMO is only observed in the cilium of a few cells in RPE1 WT cells (Figure 7A). As shown previously for the cognate dynein-2 intermediate chain, WDR60 (Vuolo et al., 2018), WDR34 KO cells show SMO localization to the cilium under basal conditions in a greater proportion of cells (Figure 7A, quantified in 7B). Expression of GFP-WDR34-WT was able to reduce basal levels of intra-ciliary SMO, albeit not to the same low level as RPE1 WT cells and not to an extent that is statistically detected as significant. p.Q158* shows a large increase in basal SMO to a level greater than in WDR34 KO cells.

**Figure 7.**
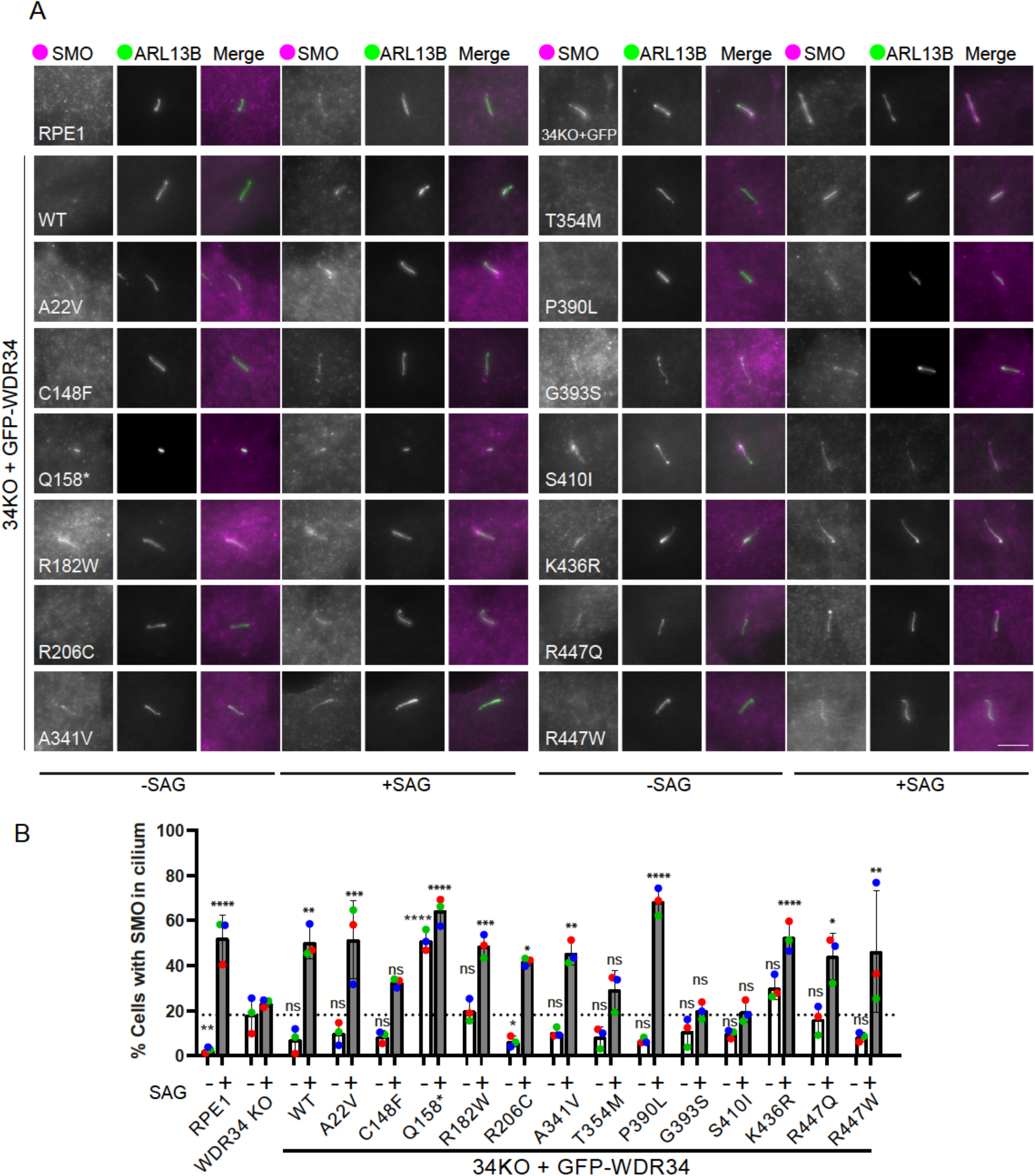
Analysis of SMO localization in ciliated cells at steady state and following SAG stimulation. (A) Cells expressing WDR34 mutations were fixed with 4% PFA followed with permeabilization by 0.1% Triton X-100 and labelled with antibodies to detect ARL13B (green) and SMO (magenta). +/-SAG stimulation are shown. Scale bar = 5 μm. (B) Quantification of these data show the basal (open bars) and SAG-stimulated (grey bars) localization of SMO to cilia. Between 28 and 117 cilia quantified in each case. Data show means and standard deviation, n=3 independent experiments. *p=<0.05, ** p<0.005, ***=p<0.001, ****=p<0.0001 (One way ANOVA with Dunnett’s multiple comparisons compared to WDR34 KO – (open bars) or + (gray bars) GFP.

SAG stimulation led to SMO entry and accumulation in cilia in WT RPE1 cells (Figure 7A, quantified in 7B). SAG treatment of cells expressing GFP resulted in only a small increase in SMO within the cilium. This contrasts with our previous work (Tsurumi et al., 2019) and (Vuolo et al., 2018) using an alternative WDR34 KO cell line and likely results from either further time in culture or the stable expression of GFP in these cells. We could clearly define that cells expressing WDR34 WT and WDR34 mutants were responsive to SAG, shown by an increased proportion of cells with SMO in the axoneme. Interestingly, p.C148F, p.G393S, p.T354M and p.S410I mutations resulted in the least SMO accumulation and are also the mutations that led to weakened assembly of the core dynein-2 proteins. This supports a key role for dynein-2 in regulating the entry of SMO as well its exit from cilia.

## Discussion

Skeletal ciliopathies caused by mutations in dynein-2 are well known for their poor correlation of genotypes and phenotypes (Huber and Cormier-Daire, 2012). Only limited work has been done on cells from these patients, in part hampered by limited availability of material. Our data provide for the opportunity to analyse the impact of many mutations in large numbers of cells. Many of the residues that are mutated are highly conserved through evolution (Supplementary Figure S3) and many are in regions likely to impact the folding or stability, either of WDR34 or of the entire dynein-2 complex (Supplementary Figure S4). We explored this using proteomics and then sought to correlate this with functional data from microscopy assays. We were unable to define any clear common outcomes in relation to complex integrity, cilia length and number, IFT, or SAG responses. These data are summarized in Figure 8. Major impacts depend, at least in part, on how effectively each WDR34 mutant is associated with other dynein-2 proteins and also likely relate to the stability of the mutant protein. Since we were unable to cluster these phenotypes, we discuss each mutant individually.

**Figure 8:**
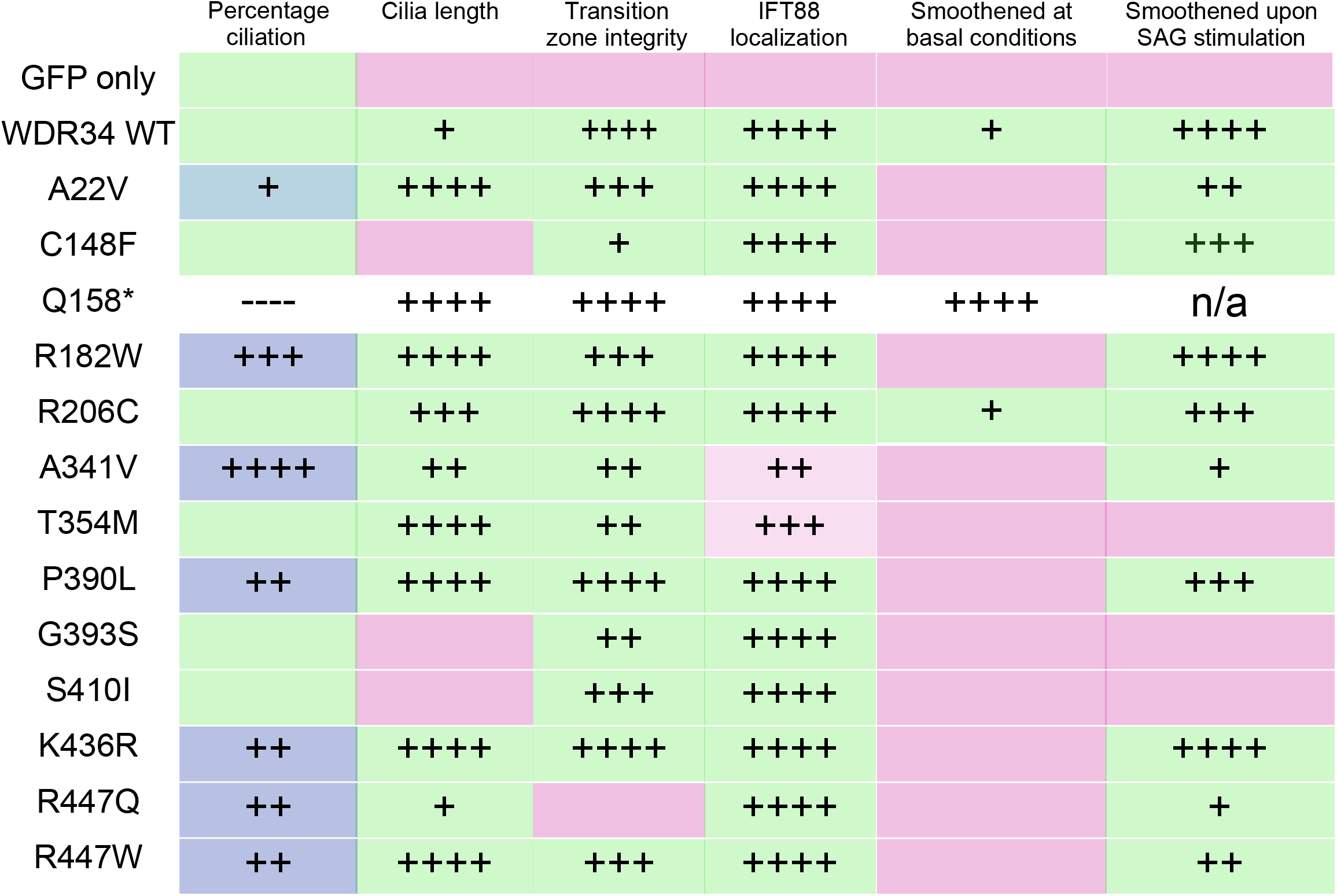
Summary of impacts of WDR34 mutations on key cilia phenotypes seen in WDR34 KO cells. Blue is used to indicate where recombinant WDR34 enhances the percentage of cells extending cilia. Green indicates restoration of function in each assay. Magenta indicates failure to restore defects. Plus signs indicate the same statistical significance shown in the corresponding main Figure. The exception is p.Q158* where outcomes are not coloured because of the dominant negative effects of this mutation. Note that A341V and T354M (light magenta) do not fully restore IFT88 localization but there are statistically detectable differences to the WDR34 KO suggesting partial rescue.

From our data, we can conclude that cilia formation can proceed in the absence of WDR34 in RPE-1 cells. Our work reconciles previously apparently conflicting findings from different KO models. We show that these changes relate to retained expression of N-terminal fragments of WDR34. In control background, p.Q158* does not impair ciliogenesis while in WDR34 KO background, p.Q158* significantly perturbs cilium formation. Our finding that the WDR34 N-terminal fragment impairs ciliogenesis in WDR34 KO background is in good agreement with Tsurumi et al. (2019). Our finding that the WDR34 N-terminal fragment does not impair ciliogenesis when expressed in a WT background (i.e. in the presence of full-length WDR34) differs from Tsurumi et al. (2019), who found that WDR34 N-terminal fragments exerted a dominant negative effect on cilia formation in control cells. This difference may relate to differences in expression level of the WDR34 N-terminal fragment; stable versus transient expression of the WDR34 N-terminal fragment (stable in our study; a mixture of stable and transient in Tsurumi et al. 2019), and the precise N-terminal fragment used (WDR34 1-157 in this study; WDR34 1-146 in Tsurumi et al. 2019). WDR34 p.Q158* was found biallelic with WDR34 p.K436R in an individual with Jeune syndrome (Schmidts et al., 2013). Our finding that WDR34 p.Q158* does not necessarily disrupt cilium formation in the background of full-length WDR34 may help to explain why WDR34 p.[Q158*]; K436R] is not embryonically lethal (as would be expected in the case of a severe cilia disruption).

p.A22V is at the N-terminal end of the protein. It has some effects on dynein-2 complex integrity, notably with regard to association with the WDR60-TCTEX module. Its expression effectively restores length and cilia number even above re-expression of WT GFP-WDR34 when expressed in WDR34 KO cells. It also fully restores the localization of RPGRIP1L and is most similar to the WT protein in terms of localization of IFT-A and IFT-B and SMO signalling. This relatively minor impact is consistent with functional predictions (Supplementary Table 2).

p.C148F shows defects in dynein-2 complex assembly or stability. It cannot restore the cilia length defect seen in WDR34 KO cells. It partially restores the steady-state distributions of RPGRIP1L and IFT-B. SMO localization in response to SAG is dampened, indicative of a defect in entry into cilia.

p.R182W results in an enhanced association with most dynein-2 subunits except for DYNLRB2. It restores cilia number even above re-expression of WT GFP-WDR34 and effectively restores cilia length, RPGRIP1L and IFT88 localization. This mutant has high levels of SMO in cilia at basal levels but retains a robust response to SAG stimulation. The SAG response remains robust indicating that the basal levels of SMO in cilia might underpin p.R182W related defects.

p.R206C shows a similar proteomic profile to p.R182W but shows other key differences. It restores the cilia length defect of WDR34 KO cells. It very effectively reverses the TZ defect shown with RPGRIP1L and IFT88 localization. A key difference to p.R182W is seen with Hh signalling where, while the SAG response also remains robust.

p.A341V also shows a similar proteomic profile to p.R182W and p.R206C with enhanced association with DYNC2H1 and WDR60 with a significant reduction in association with DYNLRB1. Its profile is very similar to p.R182W in terms of cilia number and length, RPGRIP1L localization. A key difference is seen with IFT-B localization where IFT88 remains visible throughout the cilium. The SMO profile is very similar to that seen with WT GFP-WDR34.

p.T354M shows more dramatic changes in terms of the association with other dynein-2 subunits, notably DYNC2H1, WDR60, and DYNLRB2. Notably it is also the mutation most strongly predicted to be deleterious to function (Supplementary Table 2). It restores cilia length to normal, contrasting with shorter cilia observed in patient fibroblasts (Huber et al.,2013). RPGRIP1L is restored to the base of the cilium but IFT88 persists along the entire length. The SAG-induced localization of SMO to cilia is reduced. This suggests a general reduction but not complete loss of dynein-2 activity.

Both p.A341 and p.T354 are close in proximity located on the fourth beta-propeller of WDR34 (Supplementary Figure S4). WDR34-p.A341V interaction with DYNCH1-LIC3 is increased 2-fold, whereas WDR34-p.T354M interaction with the other subcomplexes are drastically decreased. Hence, the IFT88 defect associated with the two mutations may be independent of dynein-2 assembly but depend on WDR34 folding. Both residues are located on opposite ends of consecutive beta-strands.

p.P390L associates more strongly with the DYNC2H1 and WDR60 modules with some reduction in association with DYNLRB1 and is also predicted to be deleterious to function (Supplementary Table 2). It restores cilia length and is highly effective in restoring the localization of RPGRIP1L to the base of the cilia. We see some very small defect in IFT88 localization but the SAG response mimics that of WT GFP-WDR34.

p.G393S shows the most dramatic changes in terms of association with other dynein-2 subunits with a reduction seen for all and is also that which is most strongly predicted to be deleterious to function (Supplementary Table 2). It cannot restore normal cilia length but is effective in terms of RPGRIP1L and IFT88 localization. The major defects seen with G393S are in Hh signalling with a near-complete failure to respond to SAG.

Similar to p.G393S, p.S410I fails to associate as effectively with other dynein-2 subunits and cannot restore cilia length. It restores RPGRIP1L to the base of the cilium, and IFT88 localization. It mirrors p.G393S with respect to absence of a robust SAG response. Together p.G393S and p.S410I link a failure of robust dynein-2 assembly to the ability of cells to respond to SAG; broadly, this matches the predictions of loss of function (Supplementary Table 2). WDR34 p.G393S was identified in patients with anencephaly (Yin et al., 2020). Both here and in their study, the p.G393S mutation impairs Hh signalling.

p.K436R shows enhanced association with DYNC2H1 and to a lesser extent with WDR60. Cilia number and length, IFT88 and RPGRIP1L localization are restored to wild-type phenotypes. This mutant has high levels of SMO in cilia at basal levels but retains a robust response to SAG stimulation. In this regard it is highly similar to p.R182W in terms of impacts. While p.K436R is more stable than WT WDR34 in a degradation assay (Figure 1C), p.R182W is not suggesting that phenotypes do not correlate with protein stability.

p.R447Q shows reduced association with most dynein-2 subunits. It rescues cilia number even above re-expression of WT GFP-WDR34 and retains the ability to restore cilia length, but not the localization of RPGRIP1L which remains extended into the TZ. This contrasts with shorter cilia observed in patient fibroblasts (Huber et al., 2013). However, IFT88 localization is restored indicating effective retrograde IFT. Notably, the basal levels of SMO within cilia remain high but relocalization remains responsive to SAG stimulation.

p.R447W, unlike p.R447Q, only shows a loss of association with DYNLRB1. Both changes are strongly predicted to be deleterious to function (Supplementary Table 2), yet the changes in interactions with other dynein-2 subunits do not mirror those of other mutations that are also strongly predicted to affect function (e.g. p.T354M and p.G393S). The percentage of ciliated cells is restored even above re-expression of WT GFP-WDR34 and cilia length is restored as is the localization of RPGRIP1L and IFT88. The restoration of RPGRIP1L to the ciliary base contrasts with p.R447Q. Loss of DYNLRB1 binding may underpin the severity of the p.R447W mutation (Supplementary Table 1) but the differences we observe between p.R447W and p.R447Q may alternatively be due to their different expression levels or stability where p.R447W appears more susceptible to degradation even in the presence of MG132.

Overall, there are some common themes arising from this work.

### Loss of light chain binding

We see considerable variation between the binding of dynein light chains, DYNLL1, DYNLL2 and DYNLRB1, to WDR34 mutants. Previous proteomic data showed that in the absence of WDR34, the other intermediate chain, WDR60, interacts less well with DYNLRB1 (Vuolo et al., 2018). Our data also showed that many WDR34 mutations, all located within a WD repeat domain, led to weak incorporation of the DYNLRB1 into the complex. Notably, the DYNLRB light chains lie in closest proximity to the WD repeat domains in the structure (Toropova et al., 2019). Mutations that did not affect the association of DYNLRB1 include p.A22V, which is located upstream of the DYNLRB1 binding site, p.R206C which is located between two WD repeat domains, and p.K436R in WD6. It may be that other mutations within WD repeat domains commonly affect folding or stability of the beta propeller, transmitting to the region where DYNLRB1 binds. p.K436 is close in proximity to DYNC2H1 (Supplementary Figure S2), so although the residue is in a WD repeat domain, p.K436R may instead exert its mutant effect by affecting interaction with DYNC2H1.

Using deletion mutations Tsurumi et al. (2019) showed that residues 80-93 of WDR34 are required for its interaction with DYNLL2. Our data show that the interaction is also weakened by the p.A341V mutation, again showing that mutations downstream of the region, and further along in the WD domains are also able to affect interaction with proteins at the N-terminal end. This further supports a key role for WDR34 in the incorporation light chains to the dynein-2 complex (Hamada et al., 2018; Toropova et al., 2019; Tsurumi et al., 2019; Vuolo et al., 2020).

### Impacts on IFT

In general, IFT140 localization remained consistent at the ciliary base in cells expressing GFP only or WDR34 missense mutations. IFT140 was enriched along the axoneme in cells expressing the p.Q158* truncation, similar to other models expressing short WDR34 N-terminal truncations (Tsurumi et al., 2019; Vuolo et al., 2018). IFT-A localization is affected in WDR60 KO cells (Vuolo et al., 2018) and DYNC2H1 mutant cells (Wu et al., 2017) suggesting that WDR34 may have no direct role in defining the steady-state localization of IFT-A. Localization of the IFT-B component IFT88 is abnormal in cells depleted of dynein-2 heavy chain, LIC3, WDR34, WDR60, or TCTEX1D2, or those retaining expression of short WDR34 fragments (Hamada et al., 2018; Kessler et al., 2015; Merrill et al., 2009; Taylor et al., 2015; Tsurumi et al., 2019; Vuolo et al., 2018). Expression of p.A341V and p.T354M mutations in WDR34 KO cells both failed to restore IFT88 to its normal distribution, suggesting that these residues have a more direct impact on retrograde IFT.

### Impacts on the TZ

Localization of the TZ protein RPGRIP1L is defective in WDR34 KO cells. Nearly all WDR34 mutations examined here restored this localization except for the C-terminal p.R447Q mutation. Surprisingly, p.R447Q, but not p.R447W leads to significant RPGRIP1L mis-localization, possibly due to differences in charge or steric changes impacting folding.

WDR60 is also required for RPGRIP1L localization (Vuolo et al., 2018). Since WDR34-p.R447 does not lie in the vicinity of the interface with WDR60 (Toropova et al., 2019), it may be that the intermediate chains control the TZ by different mechanisms.

### Impacts on Hh signalling

Most mutants fail to fully restore basal levels of SMO (Figure 8) to WDR34 KO cells but most partially restore function in this assay. These data require careful interpretation because of the low levels of SMO detected under basal conditions and the fact that WT GFP-WDR34 partially restores the basal localization of SMO but not to an extent that is statistically significant using ANOVA (that said, this is significant in a single t-test between only these samples). While statistical testing does not define full restoration of the normal basal level of SMO in cilia, it is evident from the images and quantification (Figure 7B, dotted line) that there is some partial restoration of function on expression of the GFP-WDR34 mutants. Hh signalling is impaired in cells depleted of dynein-2 heavy chain, WDR60, LIC3, and those retaining expression of short WDR34 fragments (Hamada et al., 2018; Taylor et al., 2015; Tsurumi et al., 2019; Vuolo et al., 2018, Wu et al., 2015). SAG-stimulated levels of SMO are impaired in some WDR34 KO cells. p.G393S and p.S410I mutations fail to restore SAG-induced translocation into cilia; p.T354M shows a partial response but this is not statistically detectable as significant. The basal level of SAG seen with the p.Q158* mutant is very high and not further increased by SAG. All other WDR34 mutants are able to restore the response to SAG.

### Perspective

Two of the mutations investigated here, p.T354M and p.R447W, were previously analysed in fibroblasts from patients with these mutations. Contrary to our results, both were found to result in shorter cilia (Huber et al., 2013). One reason for the inconsistency is that we are overexpressing the mutant proteins at much higher levels than normal, to be able to quantify large sample numbers which might restore some function.

As we were preparing this work for publication, Antony and colleagues (Antony et al., 2022) released a preprint that included analysis of two of the WDR34 mutants, p.R182W and p.G393S, included here using base editing of IMCD3 cells. This more precise approach to engineering mutants clearly has advantages in terms of engineering mutations into the endogenous locus and therefore avoiding limitations of overexpression. However, it comes with huge challenges regarding efficiency, complicated by the near-triploid state of IMCD3 cells. Their work defines more subtle, and sometimes contrasting phenotypes to ours. For both mutants, no significant changes to the ability of cells to form cilia, or in cilia length (Antony et al., 2022). Both mutants showed an increase in IFT88 at the ciliary tip, consistent with a defect in retrograde IFT. The data show a moderate increase in expression of GLI3 in both mutants as well as impaired Gli processing on SAG addition. These data are consistent with Shh signalling defects. In contrast, we find that overexpression of these mutants in a WDR34 KO background shows more clear defects with G393S showing the most significant changes in binding to other components of the dynein-2 complex and in Shh signalling. In contrast, our data show that p.R182W shows enhanced binding to dynein-2 subunits and can restore defects in ciliogenesis and IFT but not basal SMO accumulation in cilia. These differences are perhaps unsurprising given the very different model systems and in our case, we conclude that overexpression of p.R182W and p.G393S leads to more exaggerated outcomes than one would see with in locus base editing. Indeed, overexpression of p.G393S could even be acting as a dominant negative mutation. That said, expression levels of wild-type and mutant forms of WDR34 are broadly comparable in our experiments and we see robust functional rescue of many of the defects seen following KO of WDR34. Both approaches clearly have their merits. Base editing likely better reflects the clinical situation and gives a useful platform to screen for ways to alleviate phenotypes; our overexpression system provides some more insight into the biology of the system potentially leading to ways to achieve such interventions. Examples could include stabilizing assembly of the dynein-2 complex, controlling expression of mutant forms of WDR34, or targeting IFT or Hh signalling more selectively.

Patients with Jeune and short-rib polydactyly syndrome experience multi-organ abnormalities (Huber et al., 2013; Schmidts et al., 2013; You et al., 2017). Here, we have only studied a limited engineered cell system using epithelial cells. Cilia function is both cell-type and tissue-specific (Wheway et al., 2018). That said, dynein-2 is a core ciliary machine so many functions would be highly conserved. Of the mutations investigated in this study, polydactyly was reported in individuals with p.R182W, p.R206C and p.R447W mutations (Huber et al., 2013; Schmidts et al., 2013; You et al., 2017). However, in this study, none of these mutations led to an increased baseline level of SMO in cilia. This might reflect cell-type specific differences.

All *WDR34* mutations resulted in at least one cilia defect. Our results support the established roles of WDR34 in cilia extension, IFT-B maintenance, TZ integrity, and SMO signalling. Notably, they also suggest that WDR34 does not have a single role within the dynein-2 complex and likely has multiple complex functions in assembly of IFT trains, activation of dynein-2, and retrograde IFT. Some patients carry compound mutations in other dynein-2 or IFT proteins which likely intensify cilia defects and the clinical phenotype. This is the case with p.A22V, p.P390L and p.R447Q. This highlights the need and challenge of understanding dynein-2 assembly, not only in the context of the holoenzyme, but also in terms of co-assembly with IFT-A, IFT-B, and kinesin-2.

## Acknowledgements

We thank members of the Stephens, Roberts and Nakayama labs for continued discussion and help with this work; the Flow Cytometry and Wolfson Bioimaging Facilities for support and advice; Marc van der Kamp for help and advice on structural modelling and preparation of the figures. This work was funded by a JRPs-LEAD grant with UKRI-BBSRC from the Japan Society for the Promotion of Science (grant numbers JPJSJRP20181701 to K.N. and BB/S013024/1 to D.J.S) and BBSRC grant BB/S005390/1 to D.J.S. and A.R. Open access funding provided by University of Bristol.

## Competing interests

The authors declare no other competing or financial interests.

## Author contributions

Conceptualization: K.N., D.S.; Methodology: C.S., L.V., B.U., N.S., Y.K., K.H., D.S.; Validation: C.S., L.V., A.G.M., N.S., D.S.; Formal analysis: C.S., L.V., A.G.M., A.J.R., N.S., D.S.; Investigation: C.S., L.V., B.U., T.M., A.G.M., Y.K., N.S., A.J.R., K.H., D.S.; Data curation: C.S., D.S.; Writing - original draft: C.S., D.S.; Writing - review & editing: C.S., L.V., B.U., A.G.M., A.J.R., N.S., Y.K., K.N., D.S.; Visualization: C.S., D.S.; Supervision: K.N., L.V., A.J.R., N.S., D.S.; Project administration: K.N., D.S.; Funding acquisition: K.N., C.S., A.J.R., Y.K., D.S.

## Data access statement

Mass spectrometry data have been deposited to the ProteomeXchange Consortium via the PRIDE (Perez-Riverol et al., 2019) partner repository with the dataset identifiers PXD032758 and PXD025686.

## Materials and Methods

### Predictions of functional impact of mutations

To make predictions of the functional impact of WDR34 mutations using the Uniprot Q96EX3 accession, we used PolyPhen-2 v2.2.3r406 (Adzhubei et al., 2010). Sequences used were from UniProtKB/UniRef100 Release 2011_12 (14-Dec-2011); Structures from PDB/DSSP Snapshot 25-May-2021 (178229 Structures); Genes from UCSC MultiZ46Way GRCh37/hg19 (08-Oct-2009). We also ran predictions using PROVEAN (Choi et al., 2012) which uses the NCBI non-redundant database (September 2012), BLAST v2.2.24+, and CD-HIT v4.5.4 using the default cut-off of -2.5. SIFT (Sorting Intolerant From Tolerant, (Ng and Henikoff, 2003)) and Grantham (Grantham, 1974) scores are also included.

### Generation of WDR34 mutations

A plasmid encoding GFP-WDR34 was used as a template for site-directed mutagenesis performed by DC Biosciences Ltd (now Aruru Molecular Ltd, Dundee, UK). All were confirmed by Sanger sequencing. The Lenti-XTM HTX Packaging System (Clontech, Saint-Germain-en-Laye, France) was used with the appropriate plasmid (generated by DC Bioscience) to generate lentiviral particles in HEK293T cells. The resultant viral supernatant was used to transduce RPE1 cells with a p.Val24Alafs*74 mutation in WDR34 (Tsurumi et al., 2019), which were then cultured in complete medium with the addition of 5 µg/ml puromycin. Cells were sorted by fluorescence activated cell sorting using the University of Bristol Flow Cytometry Facility. Pools were selected that expressed the lowest detectable levels of GFP as verified by fluorescence microscopy. We analysed pools to avoid any differences arising from growing our individual clones.

### Cell culture

RPE1 and HEK293T cells were obtained from ATCC. WDR34 KO (WDR34 KO-1-5), derived by genome engineering from RPE1 cells are described in (Tsurumi et al., 2019). HEK293T cells were cultured in in Dulbecco’s modified Eagle’s medium (DMEM) supplemented with 10% FBS, whereas RPE1 cells were cultured in DMEM/F12 supplemented with 10% FBS. For ciliogenesis, cells were washed in PBS followed by serum starvation, by incubation in with media without FBS, for 24 h. All cells were grown at 37°C and 5% CO2.

Five of the WDR34 KO cell lines expressing GFP proteins were contaminated with mycoplasma. They were treated with plasmocin (Invivogen) for two weeks as per the manufacturer’s guidelines. All cells were from then on grown in the absence of plasmocin and verified independently to be free from mycoplasma contamination (MWG Eurofins).

To induce Hh signalling, cells were seeded, and the next day treated with 200 nM SAG, together with serum starvation for 24 hour before harvesting of cells.

### Antibodies

Antibodies used were as follows. Acetylated tubulin (Sigma-Aldrich, Cat# T6793, RRID:AB_477585, used 1:1000 for immunofluorescence, MeOH fixation); ARL13B (Proteintech, Cat# 17711–1-AP, RRID:AB_2060867, 1:1000 for immunofluorescence, MeOH fixation); ARL13B (Proteintech, Cat# 66739-1-Ig, RRID:AB_2882088, 1:1000 for immunofluorescence, MeOH or 4% PFA fixation); GAPDH (Proteintech, Cat# 60004-1-Ig, RRID:AB_2107436, 1:5000 for immunoblotting); GFP (BioLegend, Cat# 902601, RRID:AB_2565021, 1:5000 for immunoblotting, MeOH fixation); IFT88 (Proteintech, Cat# 13967-1-AP, RRID:AB_2121979, 1:300 for immunofluorescence, 4% PFA fixation); IFT140 (Proteintech, 17460-1-AP, RRID:AB_2295648, 1:100 for immunofluorescence); RPGRIP1L (Proteintech, Cat# 55160–1-AP, RRID:AB_10860269, 1:200 for immunofluorescence, MeOH fixation); Smoothened (Santa Cruz Biotechnology, Cat# sc-166685, RRID:AB_2239686, 1:100 for immunofluorescence, 4% PFA fixation).

### Immunofluorescence

Uncoated #1.5 glass coverslips were used for the culturing of cells for immunofluorescence microscopy. Depending on the antibody to be used, cells were fixed with −20°C methanol for 5 min, or with 4% PFA for 10 mins before extraction using 0.1% Triton X-100/PBS. Cells were blocked in 3% bovine serum albumin (BSA)/PBS for 30 min, followed by incubation in primary antibody solution in 3% BSA/PBS for 1 h, then in secondary antibody and DAPI (0.5 μg/ml) solution in 3% BSA/PBS for 1 h, all with PBS washes in between. Coverslips were mounted on microscope slides with Mowiol 4-88 mounting medium.

Cells were imaged using an Olympus IX-71 widefield microscope with a 63x objective, and excitation and emission filter sets (Semrock, Rochester, NY) controlled by Volocity software (version 4.3, Perkin-Elmer, Seer Green, UK). Images were processed using FIJI (Schindelin et al., 2012).

### Immunoblotting

For mutant stability assays, confluent cells were treated with +/-100 µg/ml cycloheximide +/-10 µM MG132 for 16 hours prior to lysis in RIPA buffer. Cells were lysed in RIPA buffer (50 mM Tris-HCl pH 7.5, 150 mM NaCl, 1% Triton X-100, 1% sodium deoxycholate, 0.1% SDS, 1 mM EDTA and protease inhibitors (539137, Millipore)). LDS buffer was added to clarified lysates. Protein samples were separated on 4-12% Bis-Tris gels and transferred onto nitrocellulose membranes by wet transfer. Membranes were blocked with 5% milk/PBST for 1 h, incubated in blocking solution with primary antibodies overnight, then incubated with HRP-conjugated secondary antibodies for one hour. Blots were developed by chemiluminescence.

### Immunoprecipitation

Cells ready for harvest were first incubated with 1 mM DSP for 30 min on ice to cross-link proteins. The reaction was quenched by the addition of 500 mM Tris-HCl pH 7.5 for 15 min. Cells were then washed with PBS and scraped off the plate in lysis buffer (10 mM Tris pH 7.4, 150 mM NaCl, 0.5 mM EDTA, 0.5% IGEPAL and protease inhibitors) followed by incubation on ice for 30 min. Clarified lysates were retrieved by centrifugation and incubated for 3 h on a rotating wheel at 4°C with 20 µl GFP-Trap beads. Beads were washed three times in washing buffer (10 mM Tris pH 7.4, 150 mM NaCl, 0.5 mM EDTA). Beads for proteomics analysis were resuspended in minimal volume of wash buffer.

### Proteomic analysis

Proteomic analysis including Nano LC mass spectrometry and primary data analysis was done as described in Vuolo et al (2018). Briefly, RPE-1 cells stably expressing WDR34 proteins were washed with PBS and incubated with crosslinker solution (1 mM DSP, Thermo Fisher Scientific #22585) for 30 min on ice. The reaction was quenched by adding 500 mM Tris-HCl pH 7.5 for 15 min. Immunoprecipitation of lysates of hTERT-RPE1 cells stably expressing HA-tagged WDR60 or WDR34 using anti-HA agarose beads (Sigma-Aldrich). Lysis buffer containing 50 mM Tris/HCl, pH 7.4, 1 mM EDTA, 150 mM NaCl, 1% Igepal and protease inhibitors (539137, Millipore) was used for HA immunoprecipitates and a buffer of 10 mM Tris/HCl pH7.4, 50 mM NaCl, 0.5mM EDTA, protease inhibitors and 0,5 % Igepal was used for GFP immunoprecipitates. Subsequently, cells were incubated on a rotor at 4 °C for 30 min and then lysates were centrifuged at 13,000 g at 4 °C for 10 min. Cell lysates were added to the equilibrated HA or GFP beads and incubated on a rotor at 4°C. Next, the HA beads were washed in washing buffer containing 50 mM Tris-HCl, 150 mM NaCl, 0.5 mM EDTA, Triton X-100 0.3% SDS 0.1% and GFP beads were washed in a buffer of 10 mM Tris/HCl pH7.4, 50 mM NaCl, 0.5 mM EDTA. Subsequent proteomic analysis by nano-LC MSMS using an Orbitrap Fusion Tribrid mass spectrometer (Thermo Fisher Scientific) described previously (Vuolo et al., 2018). Data relating to GFP-WDR34 interactions are from three independent experiments. These data have been deposited to the ProteomeXchange Consortium via the PRIDE (Perez-Riverol et al., 2019) partner repository with the dataset identifier PXD032758. Supplementary Table 3 gives raw and normalized abundances as well as the normalized abundance ratio of dynein-2 components, between GFP-WDR34-Gln158* and GFP-WDR34-FL cells, in two independent experiments. These data have been deposited to the ProteomeXchange Consortium via the PRIDE partner repository with the dataset identifier PXD025686.

## Notes

### Competing Interest Statement

The authors have declared no competing interest.

### Summary of Updates

Extensive new data including new Figure 1, 2C, 8. New supplemental data, notably two new supplemental figures three supplemental tables. Two new authors have been added.

