## Supplemental for "Disease-associated mutations in WDR34 lead to diverse impacts on the assembly and function of dynein-2"

### Supplementary Figure Legends

**Supplementary Figure S1** Generation of WDR34 KO RPE-1 cell line and characterization of effect of WDR34 p.Q158* on cilia formation. (A) Alignments of sequence of WDR34 KO cell line D10 determined by Sanger sequencing of genomic DNA PCR product with reference sequence. PAM is highlighted with red box. Large deletion of 410 bp is observed. (B) Translation product of WDR34 gene of WDR34 KO vs Control cell line. Red arrow indicates complete loss of exon 1 (light-chain binding sites) along with loss of START codon. (C) GFP-WDR34-p.Q158* and GFP-WDR34 (WT) stably expressed in either control, WDR34 KO D10 and WDR34 KO 1-5 background were serum-starved for 24 h to induce ciliogenesis and stained with Arl13b and acetylated tubulin antibodies. Scale bar = 10 µm.

**Supplementary Figure S2** (A) Expression of GFP-WDR34 mutants was confirmed by immunoblotting using an anti-GFP antibody with GAPDH as a loading control. Molecular weights are shown in kDa. Note that all lanes show a non-specific band at ~60 kDa. (B) Full gels from the experiment shown in Figure 2C including tubulin as an additional control.

**Supplementary Figure S3** Multiple sequence alignment of WDR34 showing the location of clinical mutations. Well-conserved residues are highlighted in blue. p.R206C is poorly conserved between species (yellow highlight). Alignments were produced using the multiple sequence alignment tool T-Coffee.

**Supplementary Figure S4** (A) Structural representation of the mutations investigated in this study in the context of the intact inactive state of the dynein-2 complex (Toropova et al., 2019, PDB 6SC2 (dynein-2, docked into subtomogram average of the anterograde IFT-B train (Jordan et al., 2018, EMDB-4303)). (B) The location of each mutation is shown and highlighted by a white circle; the original residues are shown in each case, not that arising from mutation. Figures were prepared using Pymol.

**Supplementary Figure S5** Schematic representations of proteomic data. In each case, the circle indicates a ratio of 1 between mutant and wild-type WDR34 (i.e. no impact). Those subunits lying within the circle are more tightly associated with GFP-WDR34, those outside are less tightly associated. Distances indicate the inverse of the ratios shown in Figure 1.

### Supplementary Table Legends

**Supplementary Table S1**: **Brief summary of clinical impact of *WDR34* mutations.** Descriptions are adapted from Huber et al. (2013), Schmidts et al. (2013), and You et al. (2017).

**Supplementary Table S2**: **Predictions of the functional impact of mutations in WDR34.** Scores from PROVEAN and PolyPhen2 (PPH2) are shown; we use the prediction terminology from each algorithm which are colour coded according to impact with neutral/benign in blue, possibly deleterious in yellow, and deleterious/probably deleterious in red. FPR indicates the False Positive Rate (1 − specificity at the indicated probability), TPR the True positive rate (sensitivity at the indicated probability). SIFT (Sorting Intolerant From Tolerant, (Ng and Henikoff, 2003)) and Grantham (Grantham, 1974) scores are also included.

**Supplementary Table S3**: Abundance ratios of dynein-2 subunits found in association with GFP-WDR34-FL and GFP-WDR34-p.Q158*. Data are shown from two independent experiments. The abundances were normalized to peptide counts for GFP to account for variation in expression level.

Supplementary Figure S1

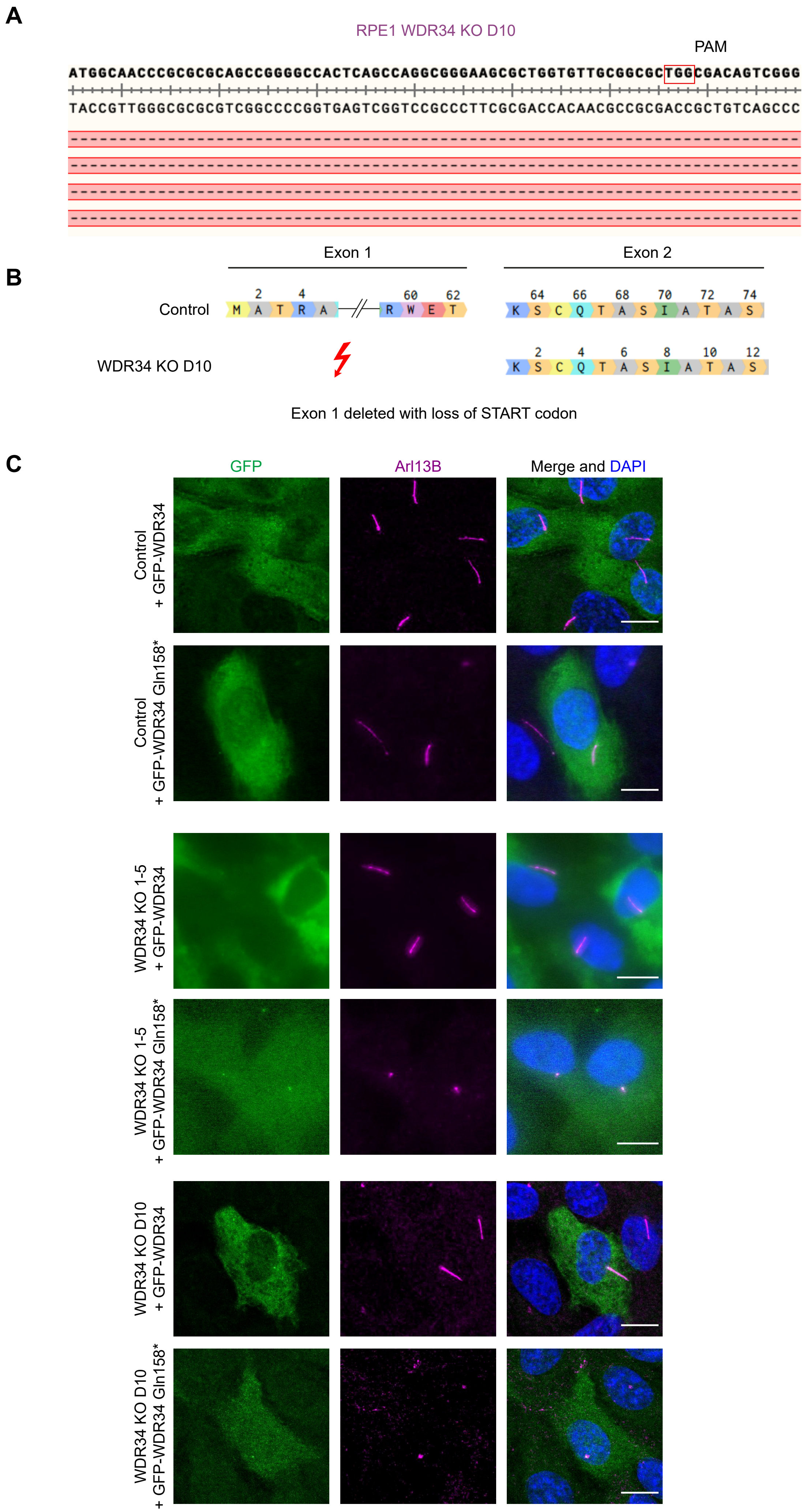

Supplementary Figure S2

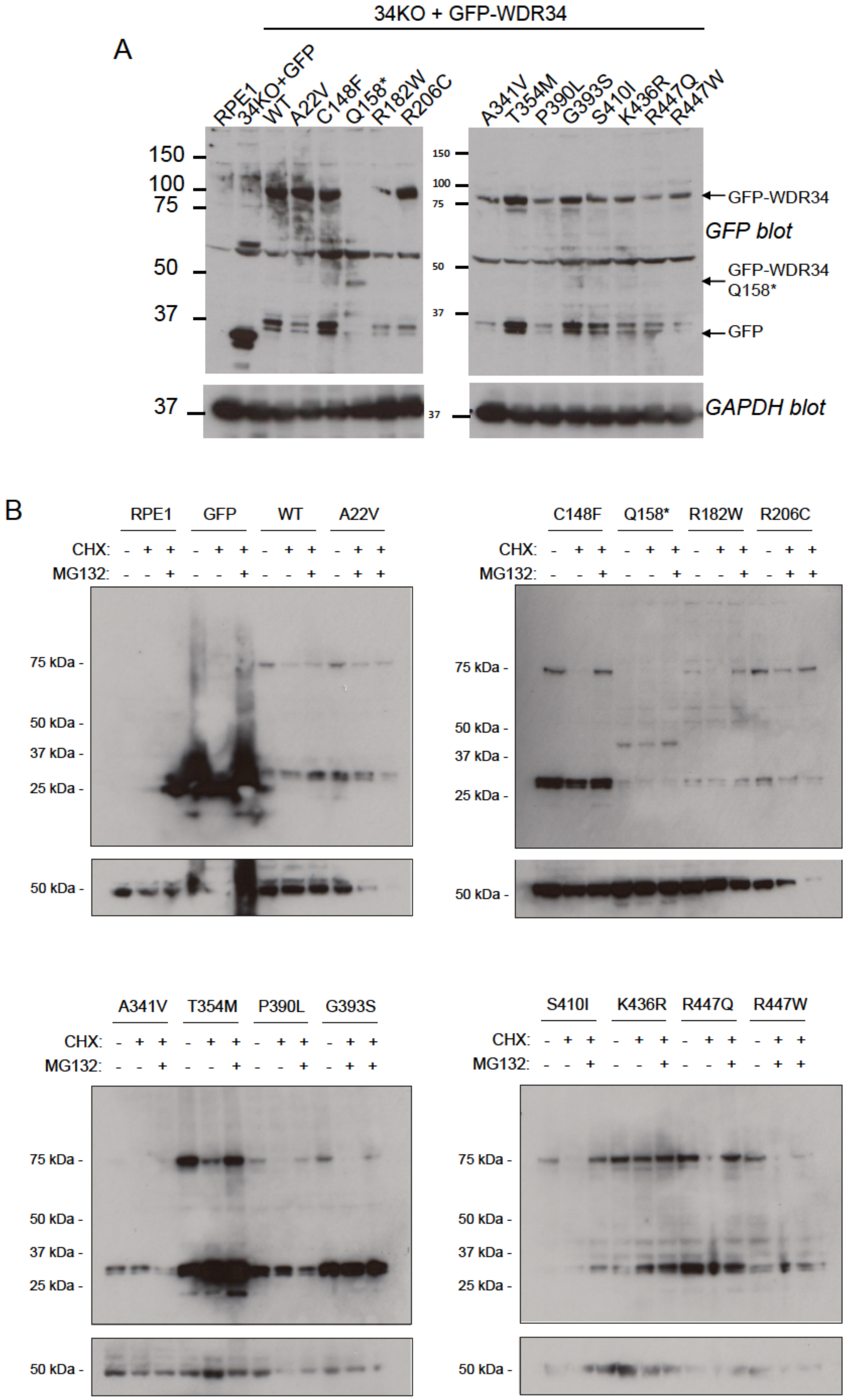

Supplementary Figure S3

| | **A22V** | **C148F** | **Q158*** | **R182W** |
|---|---|---|---|---|
| **Human** | GVAALATVGVA | QQMVSCLYTLG | GYPPAQAQGLH | ACAYGRLDHGD |
| **Chimpanzee** | GVAALATVGVA | QQMVSCLYTLG | GYPPAQAQGLH | ACAYGRLDHGD |
| **Mouse** | GAAALAAGGAG | QQTVSCLHTLV | VYPLAQGQGLH | ACAYGRLDDGD |
| **Cat** | GAEALATGGAA | QPTVTCLHTLG | GHPPAQGQGLH | ACAYGRLDDGD |
| **Dog** | GAEALATGGAA | QPTVTCLHTLG | GYPPAQGQGLH | ACAYGRLDDGD |
| **Chicken** | --SA--FTACP | NRTVLCLHTLS | SYPEAQDQHLQ | ACSYGRLNDGD |
| **Zebrafish** | ----------- | NESVSCMYRLQ | QHVDAQEKSLQ | ACGFGRVDDGD |

| | **R206C** | **A341V** | **T354M** | **P390L G393S** |
|---|---|---|---|---|
| **Human** | DRRDLRPQQPS | GATAVAFSSFD | LFILGTEGGFP | QFTFSPHGGPIYSV |
| **Chimpanzee** | DRRDLHPQQPS | GATAVAFSSFD | LFILGTEGGFP | QFTFSPHGGPIYSV |
| **Mouse** | DRQGLNPQQPS | GVTSVAFSSFD | LFVLGTEGGFP | QFTFSPHGGPVYSV |
| **Cat** | DRRGLNPQQPS | GATAVAFSGFD | LFVLGTEGGFP | RFTFSPHGGPIYSV |
| **Dog** | DRRGLNPQQPS | GATAVAFSSFD | LFVLGTEGGFP | QLIFSPHGGPIYSV |
| **Chicken** | DRRRLDPQRPD | GVTSLSFSHFD | VFIVGVEGGYS | ELAFSPHSGPLYSV |
| **Zebrafish** | DRQNLNPKRPD | GVTAVALSPWD | TFLVGSEGGLV | QFSFSPRGGPIHSV |

| | **S410I** | **K436R** | **R447Q/W** |
|---|---|---|---|
| **Human** | RNLFLSAGTDG | LQLSLKYLFAV | RWSPVRPLVFA |
| **Chimpanzee** | RNLFLSAGTDG | LQLSLKYLFAV | RWSPVRPLVFA |
| **Mouse** | RNLFLSAGTDG | LQLSHKYLFAV | RWSPVRPLVFA |
| **Cat** | RNLFLSAGTDG | LQLSHKYLFAV | RWSPVRPLVFA |
| **Dog** | RNLFLSAGTDG | LQLSHKYLFAV | RWSPVRPLVFA |
| **Chicken** | RNLFLSCGTDG | LQLSTKYLFCV | RWSPVRPLVFA |
| **Zebrafish** | RNLFVSVGTDG | LRVSDSYVFGV | RWSPTRPLVFA |

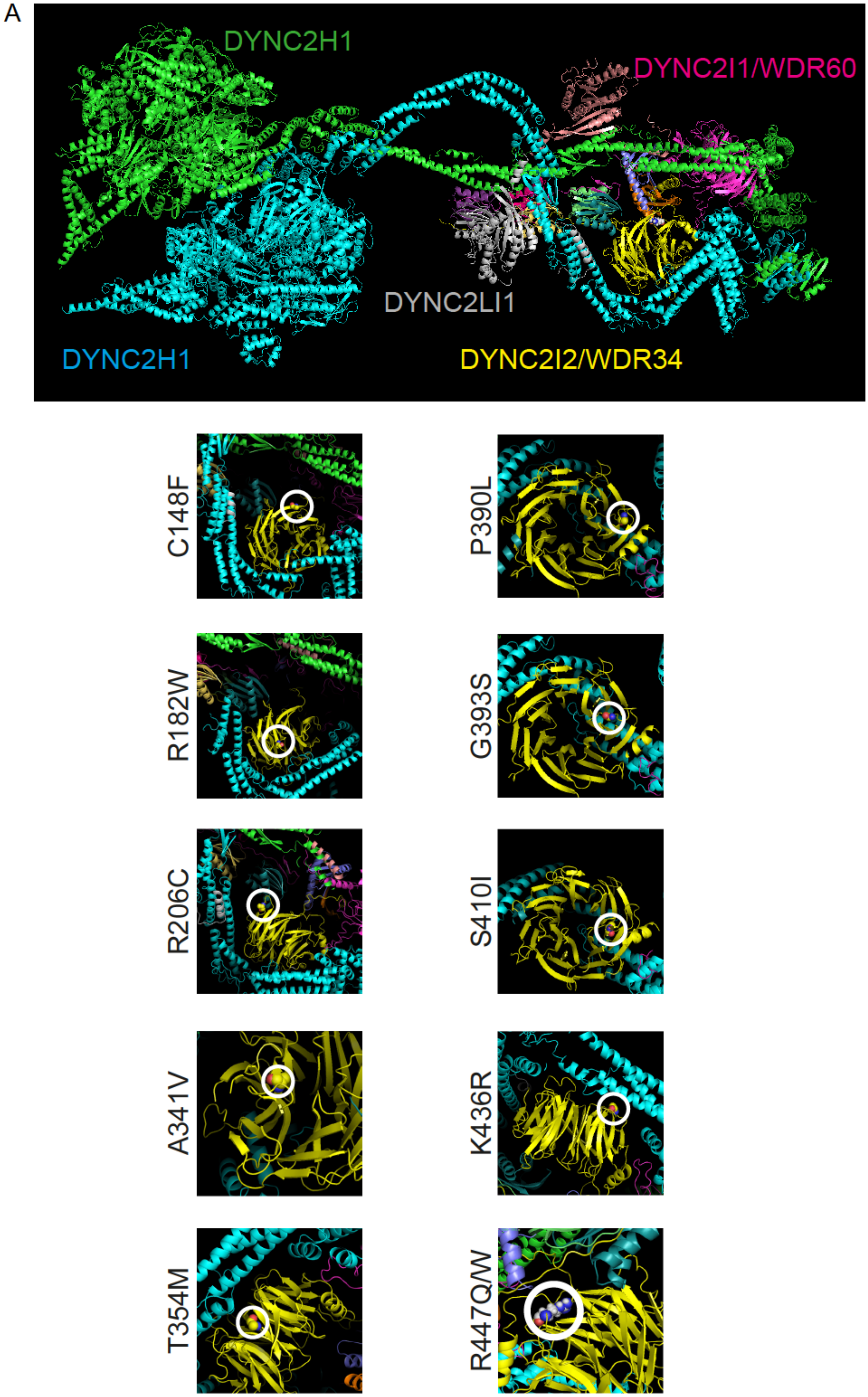
A
DYNC2H1
DYNC2I1/WDR60
DYNC2LI1
DYNC2H1
DYNC2I2/WDR34
C148F
P390L
R182W
G393S
R206C
S410I
A341V
K436R
T354M
R447Q/W

Supplementary Figure S5

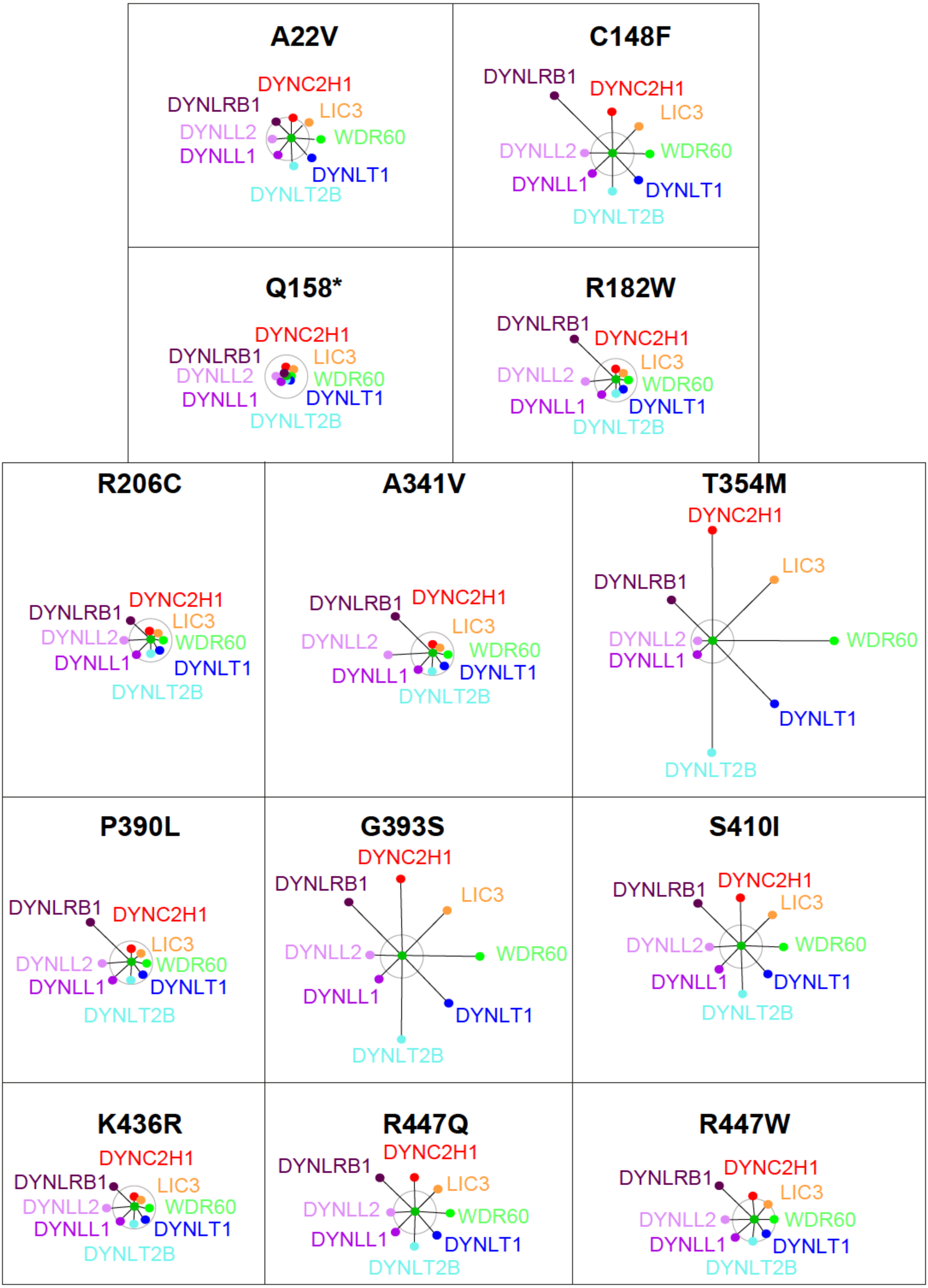

**Supplementary Table S1.**

| Mutation | Clinical impact | Comments | Reference |
|---|---|---|---|
| c.65C>T (p.A22V) | Skeletal defects | Homozygous missense identified with homozygous *DYNC2H1* and heterozygous mutations in *IFT140* c.4058C>G; | Schmidts et al., 2013 |
| c.443G>T (p.C148F) | Polyhydramnios and skeletal defects with protruding abdomen, hepatic ptosis. | Heterozygous with c.1372+1G>A | Schmidts et al., 2013 |
| c.472C>T (p.Q158*) | Skeletal defects with rod-cone dystrophy, obesity, and speech and language delay. | Heterozygous missense with c.1307A>G (K436R) | Schmidts et al., 2013 |
| c.544C>T (p.R182W) | Skeletal defects polydactyly, lung and pulmonary problems | Homozygous missense | You et al., 2017 |
| c.616C>T (p.R206C) | Skeletal defects with severe respiratory distress, recurrent infections, home ventilation. | Homozygous missense | Schmidts et al., 2013 |
| c.1169C>T (p.P390L) | Skeletal defects with bilateral nephrocalcinosis but no cysts. | Homozygous missense identified as heterozygous with *WDR19* c.2720C>T | Schmidts et al., 2013 |
| c.1022C>T (p.A341V) | Skeletal defects and, in one case, hypotrophic lungs | Homozygous missense | Huber et al., 2013 |
| c.1061C>T (p.T354M) | Skeletal defects | Homozygous missense | Huber et al., 2013 |
| c.1177G>A (p.G393S) | Skeletal defects | Missense mutation also identified as heterozygous with c.1541_1542delCA (p.T514Afs*11, is not included in this study). | Schmidts et al., 2013 |
| c.1229G>T (p.S410I) | Skeletal defects and electroretinogram at lower limit of normal | Homozygous missense | Schmidts et al., 2013 |

| | | | |
|---|---|---|---|
| c.1339C>T (p.R447W) | Skeletal defects polyhydramnios, elevated csf spaces, foot malformation, umbilical hernia, respiratory insufficiency, elevated cerebrospinal fluid spaces, foot malformation, umbilical hernia. Also identified by Huber et al 2013 in a case with skeletal defects and polydactyly. | Homozygous missense | Schmidts et al., 2013 and Huber et al., 2013. |
| c.1340G>A (p.R447Q) | Skeletal defects | Compound heterozygous missense mutation with c.982-2T>G | Huber et al., 2013 |

**Supplementary Table S1. Clinical impact of *WDR34* mutations.**

Table provides a brief description of the clinical impact of each mutation along with some contextual comments and a reference to the original descriptions.

**Supplementary Table S2**

| Mutation | PPH2 HumDiv prediction | PPH2 probability | PPH2 FPR | PPH2 TPR | PPH2 HumVar prediction | PPH2 probability | PPH2 FPR | PPH2 TPR | PROVEAN score | PROVEAN prediction (cutoff -2.5) | SIFT score | Grantham score |
|---|---|---|---|---|---|---|---|---|---|---|---|---|
| A22V | Probably damaging | 0.969 | 0.045 | 0.77 | Benign | 0.304 | 0.234 | 0.861 | -0.848 | Neutral | 0 | 64 |
| C148F | Benign | 0.251 | 0.118 | 0.911 | Benign | 0.145 | 0.29 | 0.895 | -6.533 | Deleterious | 0.06 | 205 |
| R182W | Probably damaging | 0.998 | 0.0112 | 0.273 | Possibly damaging | 0.878 | 0.109 | 0.711 | -2.362 | Neutral | 0.19 | 101 |
| R206C | Possibly damaging | 0.846 | 0.0675 | 0.834 | Benign | 0.042 | 0.382 | 0.932 | -2.738 | Deleterious | 0.03 | 180 |
| A341V | Benign | 0.34 | 0.11 | 0.9 | Benign | 0.033 | 0.4 | 0.937 | -2.137 | Neutral | 0.05 | 64 |
| T354M | Probably damaging | 0.995 | 0.0277 | 0.681 | Possibly damaging | 0.676 | 0.154 | 0.788 | -6.658 | Deleterious | 0 | 81 |
| P390L | Probably damaging | 0.962 | 0.0478 | 0.779 | Possibly damaging | 0.89 | 0.106 | 0.704 | -5.142 | Deleterious | 0.1 | 98 |
| G393S | Probably damaging | 0.999 | 0.00574 | 0.136 | Probably damaging | 0.983 | 0.0589 | 0.558 | -3.375 | Deleterious | 0.16 | 56 |
| S410I | Possibly damaging | 0.946 | 0.0537 | 0.795 | Possibly damaging | 0.583 | 0.173 | 0.81 | -0.782 | Neutral | 0.01 | 142 |
| K436R | Benign | 0 | 1 | 1 | Benign | 0.001 | 0.912 | 0.994 | -3.439 | Deleterious | 0.46 | 26 |
| R447Q | Probably damaging | 1 | 0.00026 | 0.00018 | Probably damaging | 0.994 | 0.0403 | 0.463 | -6.878 | Deleterious | 0.21 | 43 |
| R447W | Probably damaging | 1 | 0.00026 | 0.00018 | Probably damaging | 0.999 | 0.00759 | 0.0901 | -6.154 | Deleterious | 0.01 | 101 |

**Supplementary Table S2: Predictions of the functional impact of mutations in WDR34.** Scores from PROVEAN and PolyPhen2 (PPH2) are shown; we use the prediction terminology from each algorithm which are colour coded according to impact with neutral/benign in blue, possibly deleterious in yellow, and deleterious/probably deleterious in red. FPR indicates the False Positive Rate (1 – specificity at the indicated probability), TPR the True positive rate (sensitivity at the indicated probability). SIFT (Sorting Intolerant From Tolerant, (Ng and Henikoff, 2003)) and Grantham (Grantham, 1974) scores are also included.

**Supplementary Table S3.**

| NAME | Replicate 1 | | | | Replicate 2 | | | |
|---|---|---|---|---|---|---|---|---|
| | Unique Peptides | Normalised abundances | | Abundance ratio | Unique Peptides | Normalised abundances | | Abundance ratio |
| | | GFP-WDR34-Gln158* | GFP-WDR34 | GFP-WDR34-Gln158* / GFP-WDR34 | | GFP-WDR34-Gln158* | GFP-WDR34 | GFP-WDR34-Gln158* / GFP-WDR34 |
| DYNC2H1 | 163 | 17.4 | 12.6 | 1.4 | 149 | 27.2 | 14.5 | 1.9 |
| WDR60 | 36 | 9.6 | 1.7 | 5.8 | 37 | 15.7 | 2.6 | 6 |
| DYNLL1 | 2 | 2.0 | 1.0 | 1.9 | 2 | 3.1 | 1.5 | 2.1 |
| DYNLRB1 | 4 | 3.1 | 0.6 | 5.0 | 3 | 2.5 | 0.6 | 4.4 |
| DYNC2LI1 | 10 | 0.7 | 0.6 | 1.3 | 9 | 1.4 | 0.8 | 1.8 |
| DYNLL2 | 1 | 0.3 | 0.1 | 2.0 | 2 | 0.4 | 0.3 | 1.7 |
| DYNLT1 | 1 | 0.3 | 0.1 | 4.8 | 1 | 0.4 | 0.1 | 5.5 |
| TCTEX1D2 | 6 | 1.1 | 0.2 | 5.6 | 4 | 1.8 | 0.3 | 5.7 |

**Supplementary Table S3**: Abundance ratios of dynein-2 subunits found in association with GFP-WDR34-FL and GFP-WDR34-Gln158*. Data are shown from two independent experiments. The abundances were normalized to peptide counts for GFP to account for variation in expression level.
